# Multiomic profiling of human and canine soft-tissue sarcomas reveals extensive molecular homology across species and identifies clinically relevant subgroups

**DOI:** 10.64898/2026.02.19.701015

**Authors:** Daniel Fuchs, Armin Jarosch, Erin Beebe, Amiskwia Pöschel, Aaron L. Sarver, Annamaria Kauzlaric, Gustavo Ruiz Buendia, Vincent Roh, Nadine Fournier, Mikiyo Weber, Lennart Opitz, Laura Kunz, Witold Wolski, Franco Guscetti, Anne Flörcken, Mirja C. Nolff, Enni Markkanen

**Author notes:** Shared first authors (Daniel Fuchs, Armin Jarosch). Shared senior authors (Mirja C. Nolff, Enni Markkanen).

## Abstract

Soft-tissue sarcoma (STS) are rare and heterogeneous mesenchymal tumours with over 100 recognized human subtypes. Despite advances in cytogenetic and molecular characterization, diagnostic precision and therapeutic options remain limited for most subtypes. Spontaneously occurring canine STS could represent valuable translational models, due to their higher incidence and clinical similarity to human counterparts. However, molecular cross-species comparisons of specific subtypes are largely missing. Here, we performed a tissue-resolved, cross-species analysis of tumour and matched adjacent normal tissue (NT) in human and canine fibrosarcoma (FSA) and myxofibrosarcoma (MFS) by laser-capture microdissection of FFPE specimens combined with RNAseq and LC-MS/MS. Multimodal profiling revealed FSA and MFS to represent a molecular continuum rather than distinct entities in both species, resulted in identification of clinically relevant subgroups based on immune activation, proliferative activity and copy number alterations, and identified a novel canine STS subtype associated with a gene fusion. Moreover, our analyses revealed cross-species conserved transcriptomic and proteomic alterations distinguishing tumour from NT, including pathways linked to extracellular matrix remodelling, immune modulation, and cell proliferation. These data establish the first comprehensive molecular comparison of canine and human FSA and MFS, highlight the translational relevance of canine models, and identify candidate biomarkers for diagnostic refinement and development of targeted therapeutic modalities.

## Introduction

Soft-tissue sarcoma (STS) represent a heterogeneous group of rare mesenchymal tumours currently comprising over 100 different histologic and molecular subtypes in humans ^1^. For the majority of all STS entities, characteristic molecular markers and histomorphologic hallmarks are missing. The low incidence and intrinsic complexity of human STS pose diagnostic challenges in a significant number of cases and restrict understanding of disease mechanisms, contributing to poor diagnostic accuracy and suboptimal therapy ^1–3^. Subtype-specific clinical trials are very challenging due to the small number of patients, a situation that is further exacerbated by the lack of relevant animal models in which to test potential novel therapies ^4^.

Due to the closely related pathophysiology including a natural immune response, naturally occurring tumours in pet dogs present an invaluable resource for cancer research ^5,6^. Comparative study of human and canine tumours offers a unique opportunity to assess novel anti-cancer treatments in a context of naturally occurring drug resistance, metastasis and tumour-host immune interactions that are difficult to recapitulate in other models. Dogs develop morphologically and biologically comparable STS to those in humans, but much more frequently ^7–10^: STS account for 8 – 12% of all malignant dog tumours, translating to an incidence of approx. 140:100’000 ^11^, compared to less than 1% of all human adult malignancies (incidence <5:100’000) ^1,3^. Within the STS spectrum, fibrosarcoma (FSA) and myxofibrosarcoma (MFS) are a subgroup of malignant tumours of fibroblastic/myofibroblastic origin that affect both humans and dogs. Both subtypes are characterized by locally aggressive growth, low sensitivity towards radio- and chemotherapy and high recurrence rates in both species ^4,12,13^. MFS are among the most common human STS, primarily affecting elderly patients ^14–17^, and also occur frequently in dogs. In contrast, FSA are exceedingly rare in humans but among the most frequent STS in dogs ^1,11,12,18^. This positions dogs as a particularly valuable translational model, enabling structured preclinical evaluation of novel therapeutic strategies.

Although it is widely accepted that STS in dogs and humans closely share morphologic and biologic aspects, thorough molecular cross-species comparisons are currently lacking. Subtyping of human STS has evolved based on cytogenetic and molecular characterization, leading to accurate determination of various subtypes ^1^. In contrast, routine diagnosis of canine STS relies on a coarse morphology-based classification of few subtypes ^19^. This impedes direct cross-species comparisons and limits the translational utility of canine tumours as models for human STS. Moreover, therapeutic outcome in both human and canine STS patients could be greatly improved through targeted modalities, such as intraoperative tumour visualization to guide complete resection or targeted delivery of cytotoxic agents especially in the metastatic setting ^20,21^. However, development of such targeted modalities is limited by a striking lack of detailed molecular data to differentiate STS from patient-matched unaffected normal tissue (NT) for both species. While genetic, epigenetic, transcriptomic and proteomic approaches using bulk tumour tissue analysis (e.g. ^22–26^), have been applied in several human STS subtypes to identify therapeutic and prognostic targets, specific tissue-resolved proteomic and transcriptomic analyses of tumour and matched adjacent NT are not available for any STS subtype to date in either humans or dogs. Detailed insight into proteomic and transcriptomic changes between tumour and adjacent NT is urgently needed to develop more specific tools to diagnose, visualize and treat STS.

Here, we used laser-capture microdissection (LCM) followed by RNAseq and LC-MS/MS ^27–32^ to specifically analyse microscopically defined tissue areas from archival FFPE tumours. Using this approach, we performed the first simultaneous cross-species multimodal analysis of tumour and unaffected NT in human and canine FSA and MFS. The aim of this work was to perform comparative molecular characterization of STS subtypes across species and to identify molecular features that differentiate malignant tissue from the surrounding NT.

## Material and Methods

### Ethics approval and consent to participate

Human FFPE tumour tissue samples were retrieved from the archives of the Institute of Pathology of the Charité-Universitätsmedizin Berlin. The use of tissue samples derived from humans was approved by the Ethics Commission of the Charité Berlin (EA1/108/22). Canine FFPE tumour samples were retrieved from the archives of the Institute of Veterinary Pathology of the Vetsuisse-Faculty of the University of Zurich. No animals were killed for the purpose of this research project. According to the Swiss Animal Welfare Law Art. 3 c, Abs. 4 the preparation of tissues in the context of agricultural production, diagnostic or curative operations on the animal or for determining the health status of animal populations is not considered an animal experiment and, thus, does not require an animal experimentation license. The FFPE material from all canine patients was obtained for diagnostic reasons and its use for research therefore does not require ethical approval in Switzerland in accordance with the national and local legislation and institutional requirements.

### Selection of human cases for LCM

FFPE tissue samples of human STS (n = 23 FSA (comprising 3 infantile FSA, 16 adult FSA and 4 sclerosing epithelioid FSA) and n = 27 MFS) were retrieved from the archives of the Institute of Pathology of the Charité-Universitätsmedizin Berlin. All samples had been originally diagnosed and graded for clinical purposes by an expert soft tissue and bone pathologist according to WHO guidelines, and diagnosis was reviewed by a second expert pathologist (AJ) before inclusion in the study. *Table 1* and *Suppl. Fig 1A* provides clinical details for all human cases included in the study.

**Table 1.** Overview of human STS cases included in the study. AT/CT/SM indicate which matched NT were collected for the respective cases and analysis, where AT = adipose tissue, CT = connective tissue, SM = skeletal muscle, SEF = Sclerosing epithelioid fibrosarcoma, F = female, M = male.

| CaseID | Transcriptomics |  |  | Proteomics |  |  | Tumour subtype | Prior therapy | Diagnosis | Sex | Age at diagnosis | Location | Grade |
| --- | --- | --- | --- | --- | --- | --- | --- | --- | --- | --- | --- | --- | --- |
|  | AT | CT | SM | AT | CT | SM |  |  |  |  |  |  |  |
| hF01 | x | x | x | x | x | x | FSA | yes | Infantile FSA | M | 0 | Extremity | G1 |
| hF02 | - | x | - | - | x | - | FSA | yes | Infantile FSA | M | 2 | Trunk | G1 |
| hF03 | x | x | - | x | - | - | FSA | yes | Infantile FSA | F | 0 | Extremity | G1 |
| hF04 | - | - | x | - | - | x | FSA | yes | Adult FSA | M | 21 | Intraabdominal | G2 |
| hF05 | x | - | - | x | - | x | FSA | yes | Adult FSA |  |  | Pelvic | G2 |
| hF06 | - | x | x | x | - | x | FSA | no | Adult FSA | F | 33 | Extremity | G2 |
| hF07 | x | - | x | x | - | x | FSA | no | SEF | M | 51 | Trunk | G2 |
| hF08 | x | - | - | x | - | - | FSA | no | Adult FSA | F | 69 | Trunk | G1 |
| hF09 | x | - | x | x | - | x | FSA | no | Adult FSA | F | 78 | Trunk | G2 |
| hF10 | - | x | - | - | x | - | FSA | no | Adult FSA | M | 62 | Extremity | G2 |
| hF11 | - | - | x | - | - | x | FSA | no | Adult FSA | F | 69 | Extremity | G2 |
| hF12 | - | - | - | - | - | - | FSA | no | SEF | M | 25 | Extremity | G2 |
| hF13 | - | excluded | - | - | - | - | FSA | yes | Adult FSA | F | 50 | Extremity | G2 |
| hF14 | x | x | x | x | x | x | FSA | no | Adult FSA | F | 29 | Extremity | G2 |
| hF15 | - | x | x | excluded | x | x | FSA | no | Adult FSA | M | 23 | Extremity | G1 |
| hF16 | x | x | - | x | x | - | FSA | yes | Adult FSA | M | 60 | Trunk | G2 |
| hF17 | - | x | - | - | x | - | FSA | yes | Adult FSA | F | 53 | Trunk | G2 |
| hF18 | - | - | x | x | x | x | FSA | no | Adult FSA | M | 76 | Head/Neck | G2 |
| hF19 | - | - | - | - | - | - | FSA | no | SEF | M | 64 | Trunk | G3 |
| hF20 | - | - | - | - | x | - | FSA | yes | Adult FSA | F | 74 | Extremity | G3 |
| hF21 | - | - | - | x | - | - | FSA | no | Adult FSA | F | 38 | Extremity | G2 |
| hF22 | - | - | - | - | - | - | FSA | no | SEF | M | 38 | Pelvic | G2 |
| hF23 | x | - | - | x | x | - | FSA | no | Adult FSA | M | 55 | Head/Neck | G1 |
| hM01 | x | - | x | x | - | x | MFS | no | MFS | M | 78 | Trunk | G1 |
| hM02 | x | x | x | x | x | x | MFS | no | MFS | F | 56 | Extremity | G1 |
| hM03 | x | x | x | x | x | x | MFS | no | MFS | M | 52 | Trunk | G1 |
| hM04 | - | - | x | - | - | x | MFS | no | MFS | F | 87 | Extremity | G2 |
| hM05 | x | x | x | x | x | x | MFS | yes | MFS | M | 66 | Extremity | G2 |
| hM06 | x | - | x | x | - | x | MFS | no | MFS | M | 90 | Head/Neck | G2 |
| hM07 | x | x | x | x | x | x | MFS | no | MFS | M | 80 | Extremity | G2 |
| hM08 | - | excluded | - | - | excluded | - | MFS | no | MFS | M | 84 | Trunk | G1 |
| hM09 | - | x | - | - | x | - | MFS | yes | MFS | F | 52 | Extremity | G1 |
| hM10 | - | - | x | - | - | x | MFS | no | MFS | M | 87 | Extremity | G3 |
| hM11 | x | x | - | x | x | - | MFS | yes | MFS | M | 66 | Extremity | G2 |
| hM12 | x | - | - | x | - | - | MFS | no | MFS | F | 81 | Trunk | G2 |
| hM13 | x | - | x | x | - | x | MFS | yes | MFS | M | 58 | Extremity | G2 |
| hM14 | - | x | - | - | x | relabelled as CT | MFS | no | MFS | M | 59 | Extremity | G1 |
| hM15 | - | x | - | x | x | - | MFS | no | MFS | M | 60 | Extremity | G2 |
| hM16 | x | x | - | x | x | - | MFS | no | MFS | M | 84 | Extremity | G3 |
| hM17 | x | x | x | x | x | x | MFS | no | MFS | M | 61 | Extremity | G3 |
| hM18 | x | - | x | x | - | x | MFS | no | MFS | M | 88 | Extremity | G2 |
| hM19 | x | x | - | x | x | - | MFS | no | MFS | F | 96 | Extremity | G2 |
| hM20 | x | x | - | x | x | - | MFS | yes | MFS | M | 41 | Extremity | G3 |
| hM21 | x | x | - | x | x | - | MFS | yes | MFS | M | 63 | Extremity | G2 |
| hM22 | - | - | x | x | - | x | MFS | no | MFS | F | 92 | Extremity | G3 |
| hM23 | x | x | - | x | x | - | MFS | no | MFS | M | 80 | Extremity | G2 |
| hM24 | x | - | x | x | - | x | MFS | no | MFS | F | 79 | Extremity | G2 |
| hM25 | x | x | x | x | x | x | MFS | yes | MFS | F | 56 | Extremity | G2 |
| hM26 | x | - | x | x | - | x | MFS | yes | MFS | F | 74 | Extremity | G2 |
| hM27 | x | relabelled as SM | - | x | relabelled as SM | - | MFS | yes | MFS | M |  | Extremity | G1 |

### Selection of canine cases for LCM

FFPE tissue samples of canine STS (n = 30 canine FSA and n = 31 canine MFS) were retrieved from the archives of the Institute for Veterinary Pathology, University of Zurich. Cases comprised biopsies or surgical resections with the diagnosis of canine FSA or MFS and were reviewed and selected by a certified veterinary pathologist (FG) according to the criteria indicated by ^19,33^. Cases with the descriptive diagnosis Myxofibrosarcoma, which denoted tumours with FSA features displaying a substantial component of myxomatous matrix were merged into a separated group named cMFS for the purpose of this study. In addition, cMFS is herein treated as a soft-tissue sarcoma (STS) subtype, although it is not formally recognized as such in veterinary pathology ^19,33^. *Table 2* and *Suppl. Fig 1A* provides clinical details for all canine cases included in the study. Canine cases were graded based on ^34^.

**Table 2.** Overview of canine STS cases included in the study. AT/CT/SM indicate which matched NT were collected for the respective cases and analysis, where AT = adipose tissue, CT = connective tissue, SM = skeletal muscle, F = female, M = male, N = neutered, U = unknown.

| CaseID | Transcriptomics |  |  | Proteomics |  |  | Tumour Subtype | Sex | Breed | Age at diagnosis | Location | Grade |
| --- | --- | --- | --- | --- | --- | --- | --- | --- | --- | --- | --- | --- |
|  | AT | CT | SM | AT | CT | SM |  |  |  |  |  |  |
| cF01 | x | x | x | x | x | x | FSA | M/N | Dobermann | 4 | Trunk | G1 |
| cF02 | x | x | x | x | x | x | FSA | F | Bernese Mountain Dog | 10 | Trunk | G2 |
| cF03 | - | - | - | x | x | - | FSA | M/N | Great Dane | 3 | Extremity | G1 |
| cF04 | - | x | x | x | x | x | FSA | F/N | Rhodesian Ridgeback | 6 | Trunk | G1 |
| cF05 | - | x | x | x | x | x | FSA | F/N | German Shepherd | 8 | Unknown | G1 |
| cF06 | x | x | - | x | x | - | FSA | F | Rhodesian Ridgeback | 2 | Extremity | G1 |
| cF07 | x | x | - | x | x | - | FSA | M | Irish Red Setter | 6 | Extremity | G2 |
| cF08 | - | - | - | x | x | - | FSA | M/N | Boxer | 7 | Extremity | G1 |
| cF09 | x | x | x | x | x | x | FSA | M | Vizsla | 8 | Trunk | G1 |
| cF10 | - | x | x | x | x | x | FSA | M | American Staffordshire Mix | 6 | Trunk | G1 |
| cF11 | - | - | x | x | x | x | FSA | F/N | Old English B | 5 | Extremity | G2 |
| cF12 | - | x | x | x | x | x | FSA | M | Golden Retriever | 3 | Head/Neck | G2 |
| cF13 | - | - | x | x | x | x | FSA | F/N | Bichon Frisé | 11 | Head/Neck | G1 |
| cF14 | - | - | x | - | x | x | FSA | M | Beagle | 12 | Head/Neck | G1 |
| cF15 | - | - | x | - | - | x | FSA | F/N | Irish Setter | 13 | Head/Neck | G1 |
| cF16 | - | - | - | - | x | - | FSA | F/N | Pug | 6 | Head/Neck | G1 |
| cF17 | x | x | x | x | - | x | FSA | M/N | Mix | 5 | Extremity | G1 |
| cF18 | - | - | x | - | - | x | FSA | M | Briard | 5 | Head/Neck | G1 |
| cF19 | x | x | - | x | x | - | FSA | M | Akita Inu | 2 | Extremity | G2 |
| cF20 | - | x | - | - | x | - | FSA | M/N | Appenzeller Cattle Dog | 6 | Extremity | G3 |
| cF21 | - | x | - | - | x | - | FSA | F/N | Labrador Retriever | 4 | Extremity | G1 |
| cF22 | - | x | - | - | x | - | FSA | M | Jack Russel Terrier | 13 | Extremity | G2 |
| cF23 | - | - | - | - | - | - | FSA | M/N | Mix | 7 | Extremity | G2 |
| cF24 | - | - | - | x | x | - | FSA | F/N | Jack Russel Terrier | 6 | Extremity | G1 |
| cF25 | - | x | - | - | x | - | FSA | M | Hovawart | 12 | Extremity | G1 |
| cF26 | - | - | x | - | x | x | FSA | F/N | German Pinscher | 9 | Head/Neck | G1 |
| cF27 | - | - | x | - | - | x | FSA | M | Golden Retriever | 11 | Trunk | G3 |
| cF28 | - | x | x | - | x | x | FSA | F | Gordon Setter | 9 | Trunk | G1 |
| cF29 | x | x | x | x | x | x | FSA | M/N | Labrador Retriever | 11 | Trunk | G3 |
| cF30 | x | - | - | x | x | - | FSA | F/N | Mix | 11 | Perianal | G3 |
| cM01 | - | - | - | - | - | - | MFS | F/N | Golden Retriever | 10 | Trunk | G1 |
| cM02 | - | - | - | - | - | - | MFS | F/N | Giant Schnauzer | 9 | Extremity | G2 |
| cM03 | x | x | - | excluded | x | - | MFS | F/N | Groenendael | 5 | Extremity | G1 |
| cM04 | - | - | - | - | - | - | MFS | F/U | Beagle | 13 | Extremity | G1 |
| cM05 | x | x | - | excluded | x | - | MFS | M/N | Australian Shepherd | 8.5 | Extremity | G1 |
| cM06 | - | - | - | x | - | - | MFS | F/N | Appenzeller Cattle Dog | 8.5 | Trunk | G1 |
| cM07 | - | x | - | - | x | - | MFS | M/U | Malinois | 6 | Extremity | G2 |
| cM08 | - | - | - | x | - | - | MFS | F/U | Mix | 9 | Trunk | G1 |
| cM09 | - | - | - | - | - | - | MFS | F/N | Jack Russel Terrier | 11 | Extremity | G1 |
| cM10 | - | - | - | - | - | - | MFS | F/U | Mix | 10 | Trunk | G1 |
| cM11 | x | - | - | excluded | - | - | MFS | F/N | Labrador Retriever | 8 | Trunk | G1 |
| cM12 | x | - | - | excluded | - | - | MFS | F/N | Mix | 11 | Trunk | G1 |
| cM13 | x | - | - | x | - | - | MFS | M/N | Mix | 9 | Trunk | G1 |
| cM14 | - | - | - | - | x | - | MFS | M/N | Mix | 13 | Extremity | G1 |
| cM15 | - | - | - | - | - | - | MFS | U/U | Tibetan Spaniel | 7 | Extremity | G1 |
| cM16 | - | x | - | - | x | x | MFS | M/U | Miniature Schnauzer | 8 | Trunk | G1 |
| cM17 | x | - | - | x | - | - | MFS | M/N | Barbet | 7 | Extremity | G1 |
| cM18 | - | x | - | - | x | - | MFS | F | Mix | 12 | Extremity | G1 |
| cM19 | x | x | - | x | x | - | MFS | F/N | Mix | 10 | Trunk | G1 |
| cM20 | - | - | - | x | - | - | MFS | F/N | Mix | 7 | Trunk | G1 |
| cM21 | - | - | - | - | - | - | MFS | F | Malamute | 12 | Trunk | G3 |
| cM22 | - | - | - | - | - | - | MFS | F/N | Mix | - | Trunk | G1 |
| cM23 | - | - | - | x | x | - | MFS | M/N | Mix | - | Unknown | G3 |
| cM24 | - | - | - | - | - | - | MFS | F/N | Fox Terrier | - | Head/Neck | G3 |
| cM25 | - | - | x | - | - | x | MFS | F/N | Entlebucher Mountain Dog | 13 | Head/Neck | G1 |
| cM26 | - | x | - | - | x | - | MFS | M/U | Dobermann | - | Extremity | G1 |
| cM27 | - | - | - | - | - | - | MFS | M/N | Labrador Retriever | 10 | Extremity | G1 |
| cM28 | - | - | - | - | - | - | MFS | M/N | Fox Terrier | - | Head/Neck | G3 |
| cM29 | - | - | - | - | - | - | MFS | F/N | Vizsla | 11 | Extremity | G2 |
| cM30 | x | x | - | x | x | - | MFS | M/N | Mix | - | Extremity | G3 |
| cM31 | - | - | - | - | - | - | MFS | F/N | Malamute | 13 | Extremity | G2 |

### Laser-capture microdissection (LCM)

Tissue was processed for LCM as originally described ^28^ with an updated Cresyl violet staining protocol ^27^. In short, sections of 10 μm thickness mounted on PEN membrane glass slides (Thermo Scientific) were used to isolate areas of interest using the ArcturusXT^TM^ or ArcturusCellect^TM^ Laser Capture Microdissection System (Thermo Scientific) as described in ^28^. Areas to be isolated (tumour and where available matched adipose tissue (AT), connective tissue (CT) and skeletal muscle (SM)) were identified by specialist human/veterinary pathologists (AJ for human, FG for canine samples) on a consecutive 3 μm H&E-stained section, enabling detailed histomorphologic assessment of the tissue. Isolation of areas of interest was verified by microscopic examination of the LCM cap as well as the excised region after microdissection. 2 caps were collected per region of interest per case for both protein and RNA isolation, respectively. The thermoplastic membranes containing captured tissue were peeled off using sterile blades and forceps, transferred into sterile Eppendorf® Safe-Lock tubes and stored at −20°C until further processing.

### RNA isolation from FFPE tissues isolated by LCM

RNA was isolated using the Covaris® truXTRAC FFPE RNA kit and the Covaris® E220 focused ultrasonicator as established ^28–31,35,36^. RNA abundance and quality was analysed using the 4200 Tape Station Software with the High Sensitivity RNA ScreenTape kit (Agilent Technologies), according to the manufacturer’s protocol.

### RNA sequencing library preparation

10 ng of RNA diluted to a concentration of 0.33 ng/μl in a total volume of 30 μl was submitted for RNA sequencing, as detailed in ^29^. Samples not reaching these amounts/concentrations were excluded from further analysis. The SMARTer Stranded Total RNAseq Kit-Pico Input Mammalian v2 (Clontech/Takara Bio USA) was used according to manufacturer’s protocol for RNA library preparation and ribosomal RNA depletion. All samples of the same tumour type and species were processed in a single batch according to standard protocols of the Functional Genomics Centre Zurich (FGCZ). Single-end sequencing (100bp) using the Illumina Novaseq 6000 was performed for samples belonging to cFSA, hFSA and hMFS, while paired-end sequencing (150bp) using Illumina Novaseq X Plus was performed for cMFS samples together with 6 tissue samples (2 T, 2 CT, 1 AT and 1 SM) of each cohort for potential batch correction.

### RNAseq data processing

Trimming (trimgalore v0.6.7), mapping and gene counts were computed with nf-core v3.7 with option --aligner star_salmon, and --pseudo_aligner salmon (STAR v2.6.1d, salmon v1.5.2), using human (GRCh38, Gencode v94) or dog reference genome (CanFam3.1). Genes with zero TPM in all samples were filtered out. In addition, we filtered out genes with less than 20 counts on average. The gene expression was normalized between samples with the TMM method (edgeR package, v3.36.0 ^37^) and log_2_ transformed with voom (limma package, v3.50.3 ^38^). As re-sequenced samples were clustering directly next to primarily sequenced samples, we waived batch correction. A generalized linear model was used to detect differentially expressed genes incorporating adjusted (Benjamini and Hochberg method) p-values.

### Sample preparation for proteomic analysis

For proteomic analysis, all samples of the same tumour type and species were processed in a single batch, as described previously ^27^. Briefly, microdissected tissue was rehydrated by adding 900 μl of heptane and incubating for 10 min at 30°C in a thermomixer (800 rpm). After centrifugation (20’000 x g, 10 min), the heptane was removed, and the step was repeated. Subsequently, the membranes were washed with 900 μl of ethanol (5 min, RT, 1’000 rpm), 200 μl of 90% ethanol (5 min, RT, 1’000 rpm) and 200 μl of 75% ethanol (5 min, RT, 1’000 rpm). The samples were stored at -80°C overnight. The samples were then solubilized using a commercial iST Kit (Pre-Omics, Germany) with an updated version of the protocol, as described ^27^. The samples were dried to completeness and re-solubilized in 20 μl MS sample buffer (3% acetonitrile, 0.1% formic acid). Peptide concentration was determined using the Lunatic UV/Vis polychromatic spectrophotometer (Unchained Labs).

### Liquid chromatography-mass spectrometry analysis

LC-MS/MS analysis was performed on an Orbitrap Fusion Lumos (Thermo Scientific) equipped with a Digital PicoView source (New Objective) and coupled to a M-Class UPLC (Waters). Solvent composition of the two channels was 0.1% formic acid for channel A and 99.9% acetonitrile in 0.1% formic acid for channel B. Column temperature was 50°C. For each sample 1Abs of peptides were loaded on a commercial ACQUITY UPLC M-Class Symmetry C18 Trap Column (100Å, 5 µm, 180 µm x 20 mm, Waters) connected to a ACQUITY UPLC M-Class HSS T3 Column (100Å, 1.8 µm, 75 µm X 250 mm, Waters). The peptides were eluted at a flow rate of 300 nl/min. After a 3 min initial hold at 5% B, a gradient from 5 to 22 % B in 80 min and 24 to 36% B in additional 10 min was applied. The column was cleaned after the run by increasing to 95 % B and holding 95 % B for 10 min prior to re-establishing loading condition.

Samples were measured in a randomized order. For the analysis of the individual samples, the mass spectrometer was operated in data-independent mode (DIA). DIA scans covered a range from 400 to 1000 m/z in windows of 15 m/z. The resolution of the DIA windows was set to 30’000, with an AGC target value of 500’000, the maximum injection time set to 50 ms and a fixed normalized collision energy (NCE) of 33 %. Each instrument cycle was completed by a full MS scan monitoring 350 to 1500 m/z at a resolution of 120’000. The mass spectrometry proteomics data were handled using the local laboratory information management system (LIMS) ^39^.

### Data quality control and sample exclusion

After data processing, we visualized clustering of the tissue samples using PCA and unsupervised clustering for both RNA and protein data. Tissue samples of hM34 (AT, CT and T) showed low quality on both RNA and protein level and were removed for further analysis. Additionally, 5 AT of different cases were removed due to low quality. NT samples which clustered with their respective tumour samples were histopathologically reassessed to evaluate tumour cell infiltration, resulting in exclusion of 2 CT samples. Further, 2 CT samples were reclassified as SM after IHC evaluation, as annotated in *Table 1* and *Table 2*.

### LC-MS/MS data processing

The acquired raw MS data together with the prior published canine FSA raw MS data (PXD029338) ^27^ were processed using the FragPipe computational platform (version 19) ^40^ with MSFragger-DIA for direct peptide identification and DIA-NN (version 1.8.2) ^41^ for quantification. Spectra were searched against the Uniprot *Homo sapiens* reference proteome (UP000005640, release 2023_05; taxonomy 9606) and *Canis lupus familiaris* reference proteome (UP000002254, release 2023_05; taxonomy 9615). Carbamidomethylation of cysteine was set as fixed, while methionine oxidation and N-terminal protein acetylation were set as variable modifications. Enzyme specificity was set to trypsin/P, allowing a minimal peptide length of 7 amino acids and a maximum of two missed cleavages. Using the report.tsv output file generated by the DIA-NN quantification step in FragPipe, which reports precursor-level (elution group) abundances, various statistics per protein and tissue were computed, including number of quantifications, standard deviation, mean intensity, CV, the maximum, and the minimum number of peptides identified in a single sample. Finally, Uniprot protein identifiers were first converted to Ensembl and then gene names. Canine genes (CanFam3.1) were converted to human orthologues using Ensembl BioMart (release 100) ^42^.

### Proteomic data analysis

Protein log_2_ fold changes were computed based on normalized protein intensities using the r package prolfqua ^43^. The intensities were first log_2_ transformed and then z-transformed so that the sample mean and variance were equal. Next, a linear model was fitted with a single factor (tissue) to each protein, and tissue differences were estimated and tested using the model parameters. To increase the statistical power, we moderated the variance estimates using the empirical Bayes approach, which exploits the parallel structure of the high throughput experiment ^44^. Finally, the p-values were adjusted using the Benjamini and Hochberg procedure to obtain the false discovery rate (FDR).

### Immunohistochemistry

Immunohistochemical (IHC) stainings were performed on FFPE tissue sections of 1 – 2µm thickness. Sections were deparaffinized with BenchMark XT automated slide stainer (Ventana Medical Systems, Inc., AZ, USA) according to the manufacturer’s instructions. Staining was carried out with the UltraView DAB Detection Kit (Ventana Medical Systems, Inc., AZ, USA) in combination with the respective primary antibodies and haematoxylin bluing reagent. Sections were stained with H&E and primary antibodies against pan-cytokeratin (Clone AE1/AE3, ready-to-use, Dako), vimentin (clone V9, ready-to-use, Dako), smooth muscle actin (clone 1A4, 1:50, Dako) and desmin (clone D33, 1:100, Dako).

### Statistical methods

Data processing, plotting, and statistical analysis were conducted in R (version 4.3.3).

### Principal component analysis (PCA)

Principle component analysis was performed applying prcomp on normalized gene and protein intensity values. Visualization was achieved using R package ggplot2 ^45^ or scatterplot3d ^46^, with PC1, PC2 and PC3 as x, y and z axis values, respectively.

### Venn diagram

Venn diagrams were produced using either ggvenn R package ^47^ or BioVenn R package ^48^ for proportional diagrams. Identification of tumour and NT-specific proteins was performed by separating data according to tissue group and filtering by row mean !=0 to ensure presence in at least one sample. The intersection of each tissue group was used to calculate overlapping proteins and separate tissue-specific targets. Furthermore, the intersection of differentially expressed genes and proteins between tumour and the different NT was used to identify common differential expressed genes and proteins.

### Heatmaps

Heatmaps were generated using R package ComplexHeatmap ^49^. To assess high, mid and low expression in tumour samples, row clustering distance was set to “Euclidean”. Top 500 highly and lowly expressed genes and proteins were used as input for GSEA with molecular signatures database (MSigDB) ^50,51^.

### Pie charts

Pie charts were plotted using the PieChart function from the lessR R package ^52^.

### Gene set enrichment analysis (GSEA) and over-representation analysis (ORA)

To analyse functional annotation of the genes identified as differentially expressed or assigned to co-expression modules, GSEA using the web tool WebGestalt 2019 (http://www.webgestalt.org) ^53^ or molecular signatures database (MSigDB v2023.2.Hs) ^50,51^ was performed. Additional pathway analysis was performed with the help of QIAGEN Ingenuity Pathway Analysis (IPA, QIAGEN Inc., https://digitalinsights.qiagen.com/IPA) ^54^ comparing differentially expressed proteins in tumour to each NT.

### Single sample GSEA (ssGSEA)

In addition to gene set enrichment analysis, we also performed ssGSEA using the public server from GenePattern (https://www.genepattern.org/#gsc.tab=0) ^55^ to calculate separate enrichment scores for each pairing of a gene set and tumour sample.

### Gene cluster identification and GCESS calculation

To identify meaningful transcriptional patterns between tissue samples, we used the counts per million (CPM) normalized data and removed samples with less than 13’000 genes identified in human and 11’000 genes identified in canine samples. After adding a pseudocount of 1, the data was log_2_ transformed and mean-centered, and only highly variant genes (variance > 1.5) were kept. After hierarchical average linkage clustering using the Pearson similarity metric, we identified gene clusters with a dendrogram node correlation >0.6 and a minimum of 60 individual genes, calculated a GCESS score for each sample for a defined gene cluster ^56^ and performed ORA using the HALLMARK gene sets. Commonality between human and canine GCESS clusters were checked for enrichment by pairwise overlap analysis using Fisher’s Exact Test.

### Barcode plot

Cross-species comparative analysis of tumour-specific expression was performed using the barcode enrichment plot from limma ^38^, a competitive gene set enrichment analysis (cGSEA). External datasets of human sarcoma included TCGA-SARC (http://cancergenome.nih.gov/.) and GSE21122 ^57^. All target identifiers from external datasets were summarized at the gene level using BioMart ^42^. GDCquery from package TCGAbiolinks ^58^ used to obtain primary tumour RNAseq expression data for the TCGA-SARC cohort. Raw data from all datasets was log_2_ normalized and genes were ranked according to their mean expression across all samples. The barcode plot shows the ranked position of genes in one cohort (x-axis) compared to the ranked position of another dataset indicated by line extensions (y-axis). Only common genes in canine and human were included in the barcode plot analysis.

### AGDEX

Agreement of differential expression (AGDEX) analysis ^59^ was performed using our RNAseq data generated in this study to explore similarity and conservation of deregulation driven by the tumour in canine and human FSA and MFS compared to their healthy peritumoral tissue. AGDEX R package v1.54.0 was used to perform analysis with a minimum number of permutations set to 100’000. Permutation analysis was used to assess the statistical significance between species’ correlation of the differential expression results.

### Deconvolution

To estimate immune cell infiltration within the human tumour samples, we used the Timer2.0 web server (http://timer.comp-genomics.org/timer/) with our RNAseq data to receive results from different deconvolution tools: CIBERSORT ^60^, EPIC ^61^, MCP-counter ^62^, TIMER ^63^, quanTIseq ^64^, xCell ^65^. Further, we used the immunedeconv R package (v2.1.3) ^66^ to use other deconvolution tools named ConsensusTME ^67^ together with ESTIMATE ^68^ to assess scores for tumour purity together with stromal and immune abundance. Results were visualized using the R packages ggplot2 ^45^ and ComplexHeatmap ^49^.

### Identification of genomic changes

Using our generated RNAseq data we elaborated possible genomic changes using STARFusion ^69^ and SuperFreq ^70^. STAR-Fusion was used to predict fusion transcripts from RNAseq samples. For the human samples we used a pre-compiled CTAT genome LIB provided by STAR-Fusion, while for the canine samples we had to generate a custom CTAT genome Lib using the CanFam3.1 gene reference and the according dfam reference file of transposable element DNA sequence alignments ^71^. Where available, paired-end sequencing files were used. Predicted gene fusions were in silico validated using FusionInspector ^72^. Gene fusions concurrently identified in NT were filtered out.

To predict somatic mutations and copy number alterations (CNA) we used SuperFreq ^70^: We first generated BAM files using Nextflow’s standard RNAseq processing pipeline and called the variants using samtools::mpileup and varscan::mpileul2cns as suggested by the author and then fed the results to SuperFreq. As the tool relies on annotated human databases such as COSMIC or dbSNP it is unfortunately not adapted to CNA calling on canine RNAseq samples. Statistical significance of CNA was assessed using GISTIC (v2.0.23) ^73^, based on copy number segments generated by SuperFreq and the hg38.UCSC.add_miR.160920.refgene reference file. Tumour mutational burden (TMB) was estimated by dividing the number of variants called by VarScan by the size of the expressed transcriptome (defined as regions with ≥10 reads of coverage), thereby normalizing TMB per megabase of expressed sequence.

For analysis and visualisation of small nucleotide polymorphisms identified using SuperFreq, the R package maftools v2.24 ^74^ was used. Mutational signatures were called using single base substitutions (SBS v3.4) signature ^75^.

### SingleR for interspecies comparison

To evaluate how similar the canine tissue samples are to their human orthologue, we used SingleR v2.1 ^76^ for automatic annotation. As reference we took an average in expression levels for the human tissue samples with respect to AT, CT, SM, hFSA C1, hFSA C2 and hMFS; and applied this reference to the human data set for validation purposes and to the canine data set. Similarly, we used the same approach to classify the TCGA-SARC cohort using a gene signature for the 4 identified HALLMARK clusters and assessed clinical outcome.

### Regulon analysis

Transcriptional regulatory networks were inferred and analysed using the R package RTN v2.30.0 ^77,78^. For regulon reconstruction, we inserted the normalized gene expression matrix and a comprehensive list of 1612 transcription factors compiled from Lambert et al.^79^ into the transcriptional network inference algorithm. For canine tumour samples, a Benjamini Hochberg adjusted p-value <1×10^-5^ was used, while the adjusted p-value for the human samples was adjusted for sample size accordingly using tni.alpha.adjust() function provided in the RTN package (with β = 0.2) to 1×10^-4^. Following, permutation (n = 1000) and bootstrap analysis was added. Single-sample regulon activity was estimated by a two-tailed gene set enrichment analysis (GSEA-2T) to assess differences in tumours’ regulon activity profile ^80^ and visualized in a heatmap.

### Survival analysis

Disease-free survival (DFS) and overall survival (OS) were calculated from treatment start to occurrence of the event and analysed using the R packages survminer v0.4.9 ^81^ and survival v3.5-8 ^82^. Survival estimates and median survival time are reported with the corresponding 95% confidence intervals (95% CI).

## Results

### Proteomic and transcriptomic profiling of human and canine FSA, MFS and patient-matched skeletal muscle, connective and adipose tissue

To perform tissue-resolved analysis of tumour and adjacent patient-matched NT, LCM was applied to specifically isolate tumour and matched NT (SM, CT and AT, as available) from archival FFPE tissue specimens of 23 hFSA (3 infantile FSA, 16 adult FSA and 4 sclerosing epithelioid FSA; 1 case with primary and metastasis; 14 treatment-naïve), 27 hMFS (1 case with primary and metastasis; 18 treatment-naïve), 30 cFSA and 31 cMFS (all canine samples were treatment-naïve; *Fig. 1A, Suppl. data 1+2*). Isolated tissue samples were subjected to RNAseq and LC-MS/MS followed by detailed bioinformatic data analysis.

**Figure 1.**
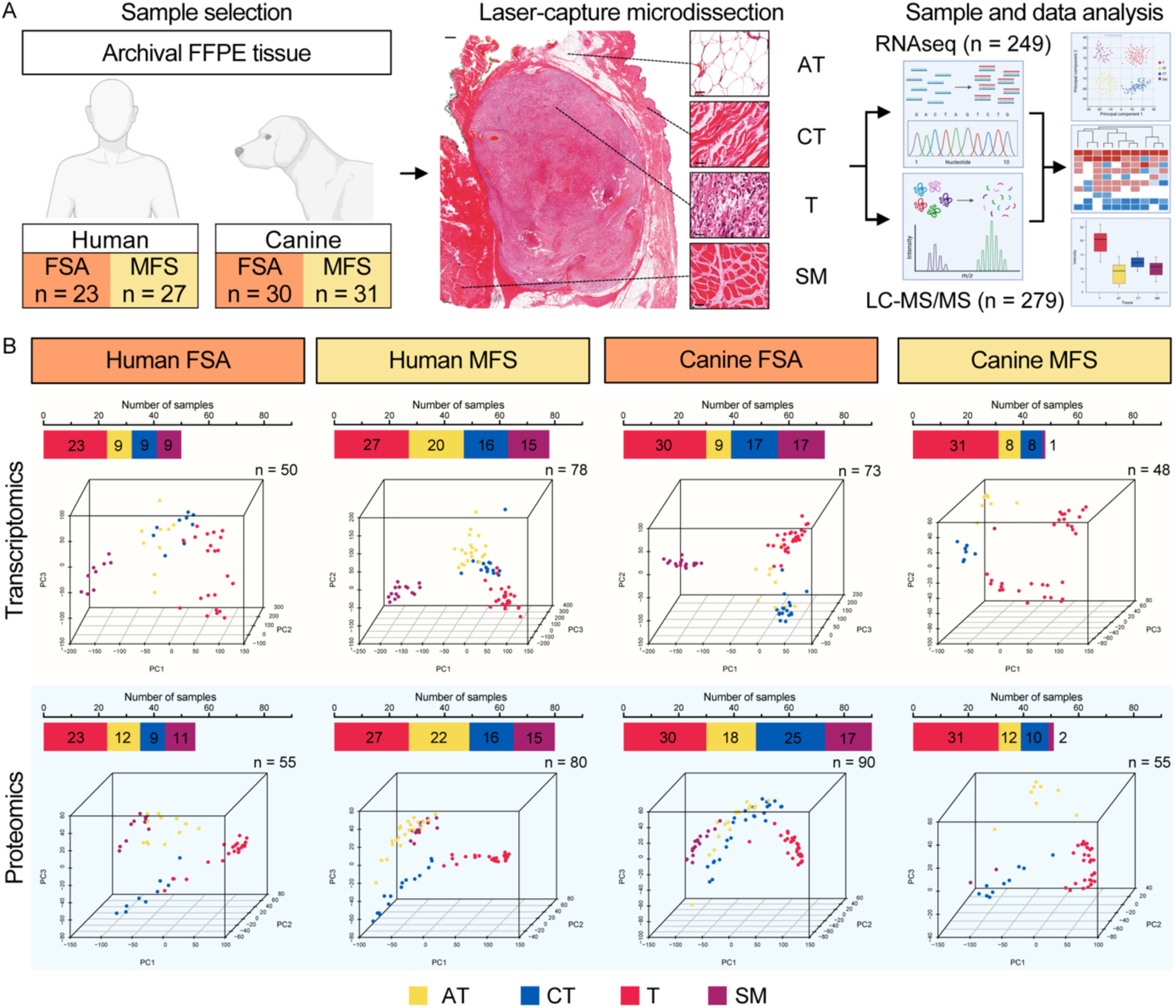
Overview of the experimental approach to characterise tumour tissue together with matched normal tissue of human and canine FSA and MFS patients. A) Laser-capture microdissection of tumour and matched normal tissue (adipose tissue, connective tissue and skeletal muscle) from FFPE sections of canine and human FSA and MFS, followed by RNAseq and LC-MS/MS (liquid chromatography tandem mass-spectrometry). The scale bars in the overview and inlets represent 1 mm and 50 µm, respectively. FSA = Fibrosarcoma, MFS = Myxofibrosarcoma. Figure created with BioRender.com. B) Number of isolated tissue samples from human FSA, human MFS, canine FSA, and canine MFS, which were submitted for RNAseq and LC-MS/MS (top) and Principal Component Analysis of samples (bottom). PCA was performed using all identified transcripts and proteins. Labels: AT = adipose tissue, CT = connective tissue, FSA = Fibrosarcoma, MFS = Myxofibrosarcoma, SM = skeletal muscle, T = tumour.

The final cohort analysed by RNAseq consists of a total of 249 specimens, including 50 samples of hFSA (23 T, 9 AT, 9 CT and 9 SM), 78 samples of hMFS (27 T, 20 AT, 16 CT and 15 SM), 73 samples of cFSA (30 T, 9 AT, 17 CT and 17 SM) as well as 48 samples of cMFS (31 T, 8 AT, 8 CT and 1 SM) (*Fig. 1B*). In hFSA and hMFS, 15’818 different transcripts were detected overall, with 11’896 and 12’965 commonly detected in all tumour samples, respectively (*Suppl. Fig. 1B).* For cFSA and cMFS, 11’751 transcripts were found overall, of which 9’446 and 9’727 were commonly shared between all tumours of the same subtype (*Suppl. Fig. 1B*).

The cohort for proteomic analysis comprises 279 specimens, including 55 samples for hFSA (23 T, 12 AT, 9 CT and 11 SM), 80 samples of hMFS (27 T, 22 AT, 16 CT and 15 SM), 90 samples of cFSA (30 T, 18 AT, 25 CT and 17 SM) as well as 55 samples of cMFS (31 T, 8 AT, 10 CT and 2 SM) (*Fig. 1B*). In hFSA and hMFS, 6’541 and 4’918 different proteins were detected overall, with 2’715 and 2’048 proteins commonly detected in all tumour samples, respectively (Supplementary Fig 1B). For cFSA and cMFS, 5’815 and 5’107 proteins were identified overall, of which 2’170 and 2’282 were commonly shared between all tumours of the same subtype (*Suppl. Fig. 1B*).

PCA of transcriptomic and proteomic data using all identified features revealed a clear separation between the four tissue types for all four STS subtypes on both RNA and protein level, supporting the validity and specificity of our analytic approach (*Fig. 1B*). Distinct sample clustering could be reproduced using unsupervised hierarchical clustering (*Suppl. Fig 2+3*).

### Molecular features in human FSA and MFS suggests these two subtypes to represent a molecular continuum of the same disease

As an entity lacking clearly defined features, we aimed to understand hFSA in more detail. Interestingly, PCA analysis and unsupervised clustering of transcriptomic data from hFSA tumour samples suggested the existence of two separate clusters within the dataset (*Fig. 1B+2A*), which was also supported by proteomics (*Suppl. Fig. 4A*). Differential expression analysis between the two clusters identified 2062 genes and 299 proteins with significant differences, of which 1362 genes and 197 proteins were highly expressed in cluster 1 (*Suppl. Fig. 4B+C*). GSEA revealed these genes and proteins to be involved in immune-related pathways and neuronal features (*Suppl. Fig. 4B+C*). While no significant difference could be observed in aspects such as sex, surgical margins, anatomical site, grading, or sample age, single sample gene set enrichment analysis (ssGSEA) confirmed a strong dichotomy between the two clusters, which was driven by 18 significantly differentially enriched gene sets involving immune and cell cycle pathways (*Fig. 2B, Suppl. Fig 4D*). This was further corroborated by deconvolution analysis and IPA revealing significant decrease in immune infiltration (adj. p-value <0.05; *Suppl. Fig. 4E*) as well as strong enrichment for expression of the immune checkpoint molecules CTLA4 and PD-L1 in hFSA C2 (*Suppl. Fig. 4F*). Of note, the cell cycle enriched cluster (hFSA C2, beige) contains a pair of strongly clustering primary tumour and metastasis (samples hF05 and hF06) from the same patient. Patients with high immune infiltration (hFSA C1, blue) showed a trend for longer disease-free survival (DFS) than those with a cell cycle signature (median DFS of 669 in C2, while median DFS (mDFS) was not reached in C1), highlighting a potential influence of the immune system in controlling FSA progression (*Suppl. Fig. 4G*). Interestingly, the hMFS cases showed a similar differentiation driven mainly by immune- and cell cycle-related gene set enrichment (*Suppl. Fig. 5*). Interactions between clinical variables and survival in the cohorts were as expected and are described in (*Suppl. Fig. 6*).

**Figure 2.**
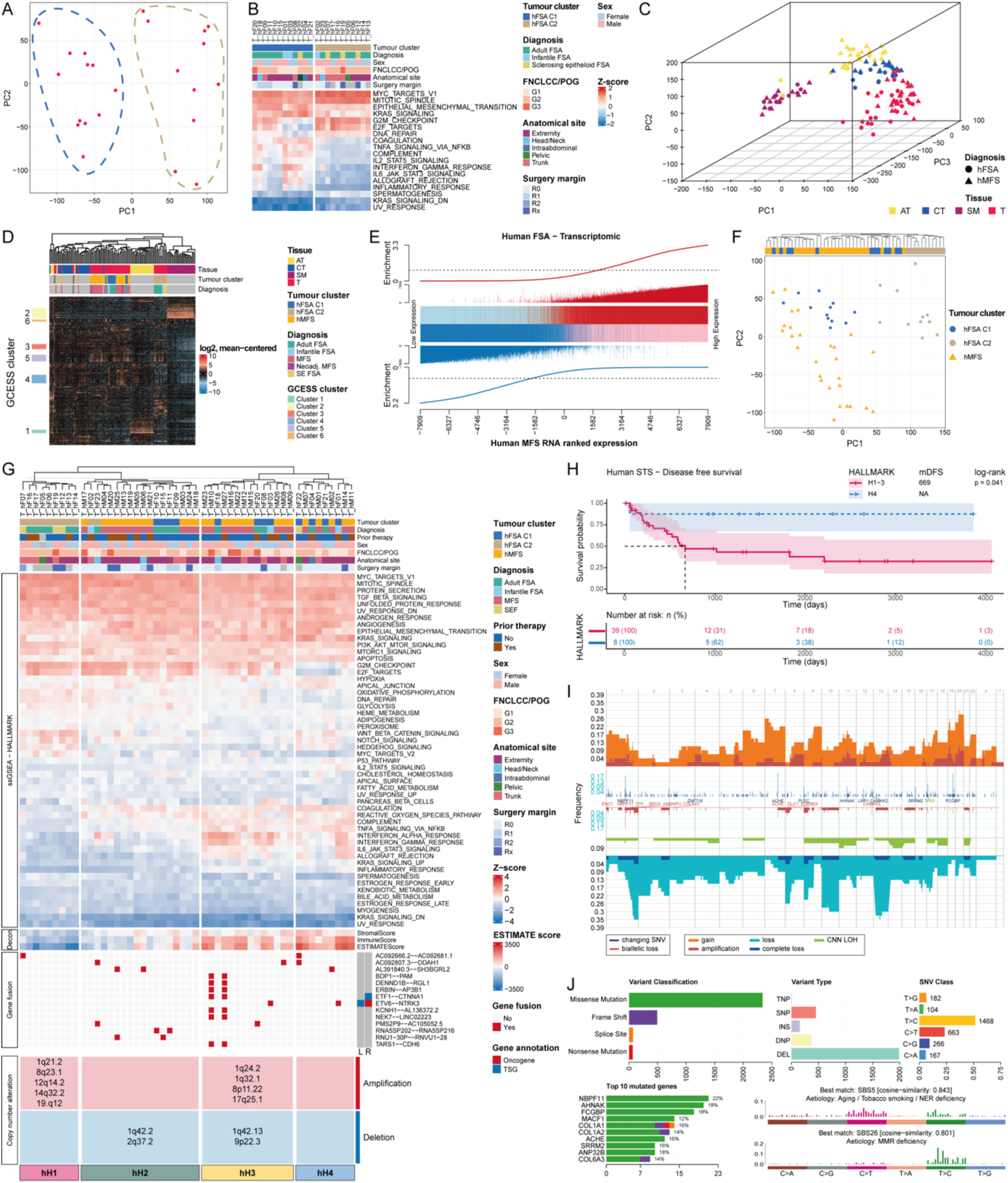
Molecular characterisation of human FSA and MFS identifies high molecular homology and clinically relevant subgroups. A) 2D PCA of 23 hFSA tumours using all identified genes. B) Heatmap showing all significant differently enriched gene sets between hFSA C1 and C2 using ssGSEA with the HALLMARK gene sets. Displayed are 18 gene sets that showed an estimated difference >0.5 or <0.5 and an adj. p-value <0.05 using student’s t-test with subsequent Bonferroni post hoc test. Clinical characteristics are annotated. C) 3D PCA of the merged hFSA and hMFS cohort using all identified genes. AT = adipose tissue, CT = connective tissue, SM = skeletal muscle and T = tumour. D) Heatmap after CPM transformation, pseudocount (+1) addition, log2 transformation, mean-centering and using highly variable genes. Gene clusters with Pearson correlation >0.6 and a minimum gene size of 60 are indicated. E) Competitive gene set testing comparing hFSA to hMFS. GSEA-like running sum statistic depicting the location of hFSA genes on a ranked list of genes in the hMFS dataset. Upregulated genes are indicated as red vertical bars, Downregulated genes are indicated as blue vertical bars. F) Hierarchical clustering and 2D PCA of the merged hFSA and hMFS cohort using all genes from tumour samples only. G) Heatmap of all 52 HALLMARK gene sets using ssGSEA for the merged cohort of hFSA and hMFS. Information on prior therapy, sex, FNCLCC/POG grade, anatomical site, surgical margin, stromal and immune score estimated by ESTIMATE, gene fusion predicted in at least 2 tumour samples, and prevalent copy number alteration in genome segments identified by GISTIC is indicated. L = left gene annotation; R = right gene annotation; TSG = tumour suppressor gene. H) Kaplan-Meyer plot comparing disease free survival between hSTS H4 and all other clusters (H1-3). I) Copy number alterations estimated from RNAseq data in hFSA and hMFS, showing frequency of changes separated by chromosome. J) Bar plots showing abundance of somatic mutations with respect to variant classification, variant type, SNV class and top 10 mutated genes, and SBS signature analysis.

As hFSA and hMFS both represent malignant fibroblastic STS, we next sought to understand how these two entities compare. To this end, we combined the transcriptomic data from hFSA and hMFS in a single PCA. In addition to demonstrating very clear clustering of all NT subtypes between the datasets, this also resulted in a single tumour cluster in which hFSA and hMFS were overlapping (*Fig. 2C*). Similarly, the samples clearly clustered according to tissue type when the most variable genes were used (*Fig. 2D*). Importantly, a very distinct clustering emerged within the tumour specimens, where FSA C2 (comprised of adult FSA, sclerosing epithelioid FSA and one infantile FSA) clearly separated from the rest of the samples. In contrast, the MFS and FSA C1 samples were much less distinguishable and arranged in an interleaving pattern of both histological subtypes. Assessment of gene cluster expression summary score (GCESS) revealed clusters exclusively enriched in adipose tissue and skeletal muscle (cluster 1 and 2), strong but variable enrichment of G2M checkpoints and E2F targets in all tumour clusters (cluster 5), and FSA C2 and a half of MFS to have very low expression of genes related to immune-related and inflammatory responses (cluster 4; *Suppl. Fig. 7A*). To further assess the level of molecular homology between the two entities, we next ranked all 15’818 tumour-derived genes shared between the two datasets according to their expression level in hMFS from low to high. We then assessed enrichment of targets in the hFSA dataset in this ranked gene list based on cGSEA. Strikingly, the 50% most highly expressed genes in the hFSA dataset (red vertical bars) were strongly enriched towards the right-hand side of the hMFS signature and hence with the highly expressed genes in hMFS (*Fig. 2E*). In contrast, the 50% lowly expressed genes from hFSA were clearly enriched towards the left-hand side, associated with transcripts lowly expressed in hMFS. This was consistent with analysis performed with the proteomic data (*Suppl. Fig. 7B).* Of note, comparing our hMFS RNA dataset with published hMFS data ^57^ obtained using a bulk tumour sequencing approach revealed a much lower correlation, likely due to contaminating non-tumour tissue present in bulk analyses as opposed to microdissected specimens (*Suppl. Fig. 7C*). Similarly, both unsupervised hierarchical clustering and PCA separated C2 tumours but failed to separate the rest of the two histological entities and resulted in a mixed pattern of hFSA C1 and hMFS instead, suggesting that hFSA and hMFS represent highly comparable entities on the molecular level, with hFSA C2 characterized by low expression of immune-related gene sets and high expression of neuronal related gene sets (*Fig. 2F, Suppl. Fig. 2A+7D-F*). Furthermore, the two entities did not differ significantly with regards to DFS or OS (*Suppl. Fig. 7G and H*). These results identify a previously unrecognized subset of FSA characterized by low immune reactivity that may be associated with worse clinical outcome (hFSA C2) and suggest the two fibroblastic STS entities FSA and MFS to represent a molecular continuum of the same disease rather than two distinct entities.

Based on this, we assessed the combined cohort in more detail. Unsupervised clustering of ssGSEA data based on RNAseq using the HALLMARK gene sets identified four main clusters: one cluster (hH1) entirely composed of hFSA C2 with strongly activated Notch, WNT-beta catenin and Hedgehog signalling, a mixed cluster (hH2) containing tumours of both entities with enrichment of coagulation and reactive oxygen species pathways, and two more clusters (hH3 and hH4) containing tumours of both entities mainly characterized by activation of immune- and coagulation-related pathways (*Fig. 2G*). In accordance with ssGSEA, deconvolution estimated a high immune score for clusters hH3 and hH4 with pronounced abundance of immune cells of myeloid lineage (*Fig. 2H, Suppl. Fig. 8*). Interestingly, cluster hH4 was the only one devoid of enrichment for the gene sets G2M checkpoint and E2F targets, which were highly enriched in the three other clusters. Strikingly, patients from the immune-high and cell cycle-low cluster hH4 showed a significantly longer DFS (mDFS not reached) compared to the other three clusters (hH1-hH3, mDFS: 669 days, p-value=0.041; *Fig. 2H, Suppl. Fig. 6N*). To validate this observation, we derived a gene signature representing the four HALLMARK clusters for application to external datasets. When applied to our own cohort, the signature demonstrated high predictive accuracy, with only a single sample misclassified (*Supplementary Fig. 9A*). Analysis of the TCGA-SARC cohort, comprising 213 soft tissue sarcoma cases across multiple histological subtypes, revealed that, in addition to an association with older age, patients classified within the hH4 cluster exhibited significantly improved overall survival and a reduced hazard ratio (*Supplementary Fig. 9*).

Hence, despite diverging histomorphology, hFSA and hMFS represent highly comparable entities on the molecular level that can be divided into four clinically relevant subgroups based on cell proliferation and immune response pathway activation status that stratify with respect to patient outcome.

### Genomic alterations as underlying driver of hFSA and hMFS behaviour

As expected, the overall somatic mutational burden was low with <2.5 Mut/Mb (average 1.19 per MB) across all human STS cases (*Suppl. Fig. 10A*) ^22^. STAR-Fusion-based prediction of gene fusions identified 78 different fusions exclusively in tumour tissue, including the gene fusion ETV6—NTRK3 in 2 out of 3 infantile FSA, all of which were FISH-positive (*Fig. 2G, Suppl. Fig. 10A-C*). Also, a similar pattern of gene fusions was identified for the two hMFS hM10 and hM27, where the second case represents the metastasis of the primary tumour and shows additional gene fusions not predicted in the primary tumour.

A wide range of CNA was estimated across all cases with a third of samples displaying a loss in chromosome 1, 10 and 13, and a fourth of samples with an estimated gain in chromosome 7 and 20 (*Fig. 2I*). GISTIC analysis further identified regions of the genome which are significantly altered within tumour clusters (FDR <0.25). Interestingly, while hFSA C1 showed no significant CNA, we identified significant amplification peaks in hFSA C2 around 1q21.1, 8q23.1, 12q14.2, 14q32.2 and 19q12, and significant deletion peaks in hMFS around 1q42.2, 2q37.2, 9p22.3 and 11q23.3 (FDR < 0.25; *Suppl. Fig 10E*). In addition, we checked for significant enrichment of CNA in the different HALLMARK clusters (*Suppl. Fig. 10D*). As H1 consists only of tumours belonging to hFSA C2, we observed the same amplification peaks in this cluster, including the region centered around 8q23.1 involving the oncogene *MYC*, an important regulator of cell proliferation, and 12q14.2 involving the oncogene *MDM2*, an important negative regulator of p53. Cluster hH2 showed only enrichment in deletions with peaks around 1q42.2 and 2q37.2. Interestingly, cluster hH3 showed significant amplification peaks around 1q24.2, 1q32.1, 8p11.22 and 17q25.1, involving *DDR2* an oncogene and tyrosine kinase receptor involved in fibroblast migration and proliferation during wound healing, and significant deletion peaks around 1q42.13 and 9p22.3 containing cell cycle regulators *CDKN2A* and *CDKN2B*. No significantly altered regions were observed for cluster hH4.

Analysis of somatic mutations revealed missense mutations and deletions as the most frequent alterations in hFSA and hMFS. Notably, recurrent mutations were detected in *NBPF11*, *AHNAK*, and *FCGBP* in approximately 20% of samples. Furthermore, mutational signature analysis indicated enrichment of processes associated with aging/tobacco smoking/ NER deficiency and MMR deficiency (*Fig. 2J*).

In summary, the two identified hFSA clusters can be distinguished based on the levels of CNA that influence the clinical patient outcome. Moreover, while tumours within cluster hH4 that is associated with a favourable clinical outcome may harbour fusion genes, they do not exhibit evidence of copy number alterations.

### Molecular assessment of canine FSA and MFS suggests these two subtypes to represent a molecular continuum of the same disease and reveals a novel canine STS subtype

With STS subtypes in veterinary medicine poorly characterized on a molecular level, differentiation of canine FSA and MFS is based on histomorphology and lacks clearly defining features. Like for the human data, PCA and unsupervised hierarchical clustering on the merged RNA data of cFSA and cMFS separated tumours from NT into one big mixed cluster of entities, indicating a strong molecular overlap between these entities (*Fig. 3A, Suppl. Fig. 3*). Interestingly, however, while the majority of cFSA (orange circle) and cMFS (yellow triangle) formed a continuous overlapping spectrum similar to the human samples, 15 tumours (14 cMFS, 1 cFSA) separated into a clearly distinct cluster (*Fig. 3B*; purple; henceforth referred to as ‘cSubcluster’), which could also be observed on the protein level (*Fig. 1B+3C*), suggesting these tumours to represent a different entity. This was further supported by using the most variable genes, which clearly separated cSubcluster, while the other samples formed a mixed population (*Fig. 3D*). Assessment of GCESS clusters revealed cSubcluster to have very low expression of genes related to immune-related and inflammatory responses (cluster 4) and high expression of epithelial-to-mesenchymal transition and angiogenesis (cluster 6 and 7), while G2M checkpoint and E2F targets were variable across all tumour samples (cluster 8). Similar to the human side, the NT showed tissue specific enrichment of oestrogen response, adipogenesis and myogenesis, respectively (cluster 2, 3 and 5; *Suppl. Fig 11A*). Indeed, 2305 genes and 391 proteins were identified as differentially expressed in cSubcluster (abs(log_2_(FC)) >1 & adj. p-value/FDR <0.05), supporting the notion that cSubcluster represents a distinct, yet undefined canine STS entity (*Suppl. Fig. 11B+C*). Further, PC loadings highlight nerval development, angiogenesis and system development as difference driver in cSubcluster (*Suppl. Fig 11D*). In contrast, direct comparison of cFSA and cMFS (without cSubcluster) using cGSEA revealed a high degree of molecular homology between the two entities on both RNA and protein level, suggesting these two subtypes to represent a molecular continuum of the same disease (*Fig. 3E, Suppl. Fig. 11E*).

**Figure 3.**
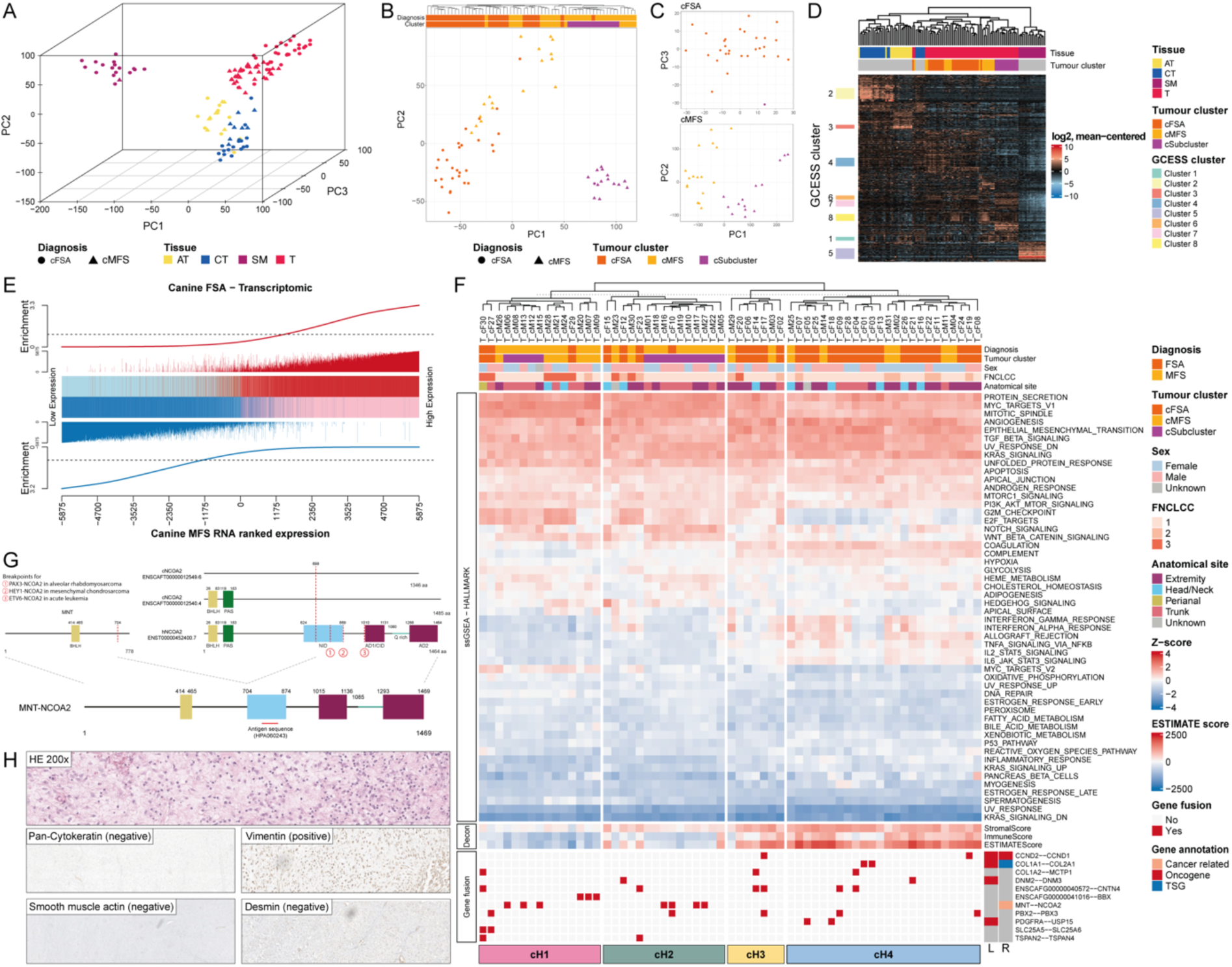
Molecular characterisation of canine FSA and MFS reveals strong molecular homology and a novel STS subtype. A) 3D PCA of the merged cFSA and cMFS cohort using all identified genes. AT = adipose tissue, CT = connective tissue, SM = skeletal muscle and T = tumour. B) 2D PCA and unsupervised hierarchical clustering of the cFSA and cMFS tumours using all identified genes. C) 2D PCA of the proteomic datasets of cFSA and cMFS displaying only tumour samples. Proteomic datasets were not merged to avoid loss of information due to batch correction. D) Heatmap of all RNA samples after CPM transformation, adding a pseudocount (+1), log2 transformation, mean-centering and using only highly variable genes. GCESS clusters with Pearson correlation >0.6 and a minimum gene size of 60 are indicated. E) Competitive gene set testing comparing cFSA with cMFS. GSEA-like running sum statistic depicting the location of cFSA genes on a ranked list of genes in the cMFS dataset. Upregulated genes are indicated as red vertical bars, Downregulated genes are indicated as blue vertical bars. F) Heatmap of all 52 HALLMARK gene sets using ssGSEA for the merged cohort of cFSA, cMFS and cSubcluster. Information on sex, FNCLCC grade, anatomical site, stromal and immune score estimated by ESTIMATE and gene fusion predicted in at least 2 tumour samples is indicated. L = left gene annotation; R = right gene annotation; TSG = tumour suppressor gene. G) Displayed are MNT and NCOA2 transcripts with respective functional domains and estimated breakpoint of MNT-NCOA2 as well as other reported breakpoints in NCOA2 in human cancers. H) H&E and IHC labelling of one representative case in the cSubcluster using markers for pan-cytokeratin, vimentin, smooth muscle actin and desmin.

To assess cFSA, cMFS and the newly identified cSubcluster more in detail, we performed ssGSEA analysis based on RNAseq data using the HALLMARK gene sets (*Fig. 3F*). While majority of the canine cases were low grade tumours, this revealed 4 clusters independent of diagnosis, similar to the human data: one cluster (cH1) composed of all three entities with strongly activated MYC targets V2, Notch, WNT-beta catenin, Hedgehog signalling, a second mixed cluster (cH2) containing mostly tumours of cSubcluster with similar enrichment profile as cH1 minus enrichment in MYC targets V2, and two more clusters (cH3 and cH4) containing tumours with high activation of coagulation- and complement system and immune-related pathways, including interferon alpha and gamma response and interleukin signalling. In line with this, deconvolution estimated a high immune score for immune-enriched clusters cH3 and cH4 (*Fig. 3F*). Interestingly, in contrast to cH1-3, cluster cH4 was the only cluster which showed no enrichment for the gene sets G2M checkpoint and E2F targets. Due to limited follow-up data, no associations with outcome could be reliably assessed in the canine cohort.

After keeping only tumour-unique gene fusions not predicted in NT samples, 44 gene fusions were identified in canine STS (*Suppl. Fig. 12A and B*). Several fusions were predicted in more than 3 cases: ENSCAFG00000040572--CNTN4 (6/29 cFSA), PBX2--PBX3 (4/29 cFSA and 1 cSubcluster case histopathology-based diagnosed as cFSA), COL1A2--MCTP1 (3/29 cFSA), ENSCAFG00000041016--BBX (3/17 cMFS), and ENSCAFG00000043897--SEMA6A (3/17 cMFS) (*Fig. 3F, Suppl. Fig. 12B*), representing gene fusions with potential for driving tumorigenesis in canine FSA and MFS. Interestingly, this identified a predicted novel chimeric MNT-NCOA2 gene fusion in 7 out of 15 tumour samples exclusively belonging to the cSubcluster (*Fig. 3F, Suppl. Fig. 12B*). Both, MNT and NCOA2 map to the reverse strand on chromosome 9 and 29 on the canine genome, respectively, indicating the involvement of a break followed by a chromosomal translocation for the development of the predicted in-frame MNT-NCOA2 gene fusion (*Suppl. Fig. 12C*). The predicted fusion protein harbours an N-terminal basic helix-loop-helix (bHLH) DNA binding/protein dimerization domain from MNT and C-terminally a disrupted nuclear interacting domain (NID) and two transcriptional activation domains (AD1/CID and AD2) provided by NCOA2 (*Fig. 3G*). Consequently, it is reasonable to assume that the chimeric protein employs the bHLH domain of MNT to heterodimerize with MAX; however, rather than repressing MYC target gene expression, it likely activates these targets via the transcriptional activation domains of NCOA2, ultimately resulting in increased cellular proliferation and survival. Indeed, our data clearly supports high activation level of MYC targets across STS subtypes in general, and in cSubcluster in particular (*Suppl. Fig. 12D*). Further, as a result of high activation of MYC targets, a negative feedback loop is highly probable to control the dysregulated gene expression by upregulating MNT expression (MYC antagonist). Indeed, we can see a significant upregulation of both MNT and NCOA2 expression to a very similar level for both genes in cSubcluster compared to cFSA and cMFS as well as NT (log_2_(FC) > 1 & adj. p-value <0.001; *Suppl. Fig. 12E*), further suggesting that these two genes are co-expressed as a fusion gene. Renewed assessment of histomorphologic features combined with IHC defined the tumours as fibroblastic lesions in proximity to the fascia with low proliferation and inflammatory infiltration. Negative staining for pan-cytokeratin and positive staining for vimentin supported a mesenchymal origin, while negative staining for smooth muscle actin and desmin excluded a muscular differentiation. Overall, these findings indicate a fibroblastic tumour with low proliferation and inflammatory infiltration. (*Fig. 3H, Suppl. Fig. 13*). Based on the histomorphologic presentation and the involvement of NCOA2-gene rearrangements in 4 human STS subtypes (mesenchymal chondrosarcoma, alveolar rhabdomyosarcoma, vulvovaginal myxoid epithelioid tumour and angiofibroma) ^83–88^, the cSubcluster shares features with human angiofibroma, a rare histologically benign but locally invasive STS subtype thus far mainly described in the nasal cavity of dogs ^89^. The prediction of the gene fusions and its potential role in MYC target expression may indicate a more malignant form. This clearly underscores the limited scope of STS classification in veterinary medicine and urges further molecular characterization in diagnosis of canine STS.

As such, we find cFSA and cMFS to represent highly comparable entities on the molecular level and identify a novel canine STS subtype characterized by a specific MNT-NCOA2 fusion.

### Extensive cross-species homology of human and canine FSA and MFS on the level of gene and protein expression

To assess whether canine STS could indeed serve as reliable model for studying human STS, we performed comparative analyses across both entities and species. To this end, we generated transcriptional signatures for tissue types in our human dataset (hFSA C1, hFSA C2, hMFS, AT, CT, SM) and used these to assign transcriptional class labels to each of the canine tissue samples (*Fig. 4A, Suppl. Fig. 14*). This classified the canine tumours diversely as hFSA C1, C2 and hMFS, suggesting the presence of the same molecular subgroups within canine FSA and MFS as in the human counterpart. List similarity in the identified GCESS clusters indicated that proliferative (hGCESS 5 vs cGCESS 8) and immune (hGCESS 4 vs cGCESS 4) clusters were defined by the same genes and could independently be observed in each dataset, further corroborating the similarity (*Suppl. Fig. 14A*). Further, PCA using the 9’685 shared transcripts revealed the different tissues to cluster in a highly similar fashion independent of the species, with the species effect only evident on PC2 (*Suppl. Fig. 14B*). Additionally, analysis of canonical pathways using IPA predicted translation and EIF2 signalling as commonly activated and extracellular matrix organization, collagen assembly, chain trimerization and degradation and complement cascade to be deactivated in tumour versus NT in both canine and human FSA and MFS (*Fig 4C, Suppl. Fig. 14C-E).* Further, we identified MYC and KRAS as repeatedly activated top upstream regulators across both species (*Suppl. Fig. 14C-E*). Remarkably, juxtaposing both hFSA and hMFS to the canine counterparts resulted in highly significant enrichment of highly and lowly expressed genes and proteins, respectively (*Fig. 4C+E, Suppl. Fig. 14I+J*). Furthermore, as expected from the molecular homology between the subtypes within each species (*Figs. 2+3*), the enrichment was conserved also in the comparison across species and subtypes on RNA and protein level, juxtaposing hFSA to cMFS (*Suppl. Fig. 14K+L*) and hMFS to cFSA (*Suppl. Fig. 14M+N*). The conservation in target expression between the species was further corroborated using agreement of differential expression for cross-species genomics (AGDEX) on the transcriptomic level (*Fig. 4D+F*). The positive values observed for both the cosine of the angle (cos) and the difference of proportions (dop) statistics suggest that the canine tumours closely recapitulate the human disease. This hypothesis is further supported by permutation analysis indicating a significant enrichment of differentially expressed genes shared between canine and human STS. GSEA analysis of the commonly up- and downregulated genes identified using AGDEX highlighted common positive enrichment for cell cycle, chromosome organization and DNA repair and negative enrichment for immune response and muscle development in both canine and human STS (*Suppl. Fig. 14O+P*). In addition, gene set enrichment analysis of differentially expressed genes in tumour versus CT using the HALLMARK gene sets highlighted further shared patterns (*Fig. 4G*) such as E2F targets, G2M checkpoints, mitotic spindle and MYC targets V1 as commonly significantly enriched gene sets, similar to the observations using ssGSEA (*Fig. 2G+3F*). Correlation analysis of the activation status in HALLMARK gene sets further revealed strong similarity between human and canine STS indicated by a high Pearson coefficient and strong p-value (*Suppl. Fig. 14Q*).

**Figure 4.**
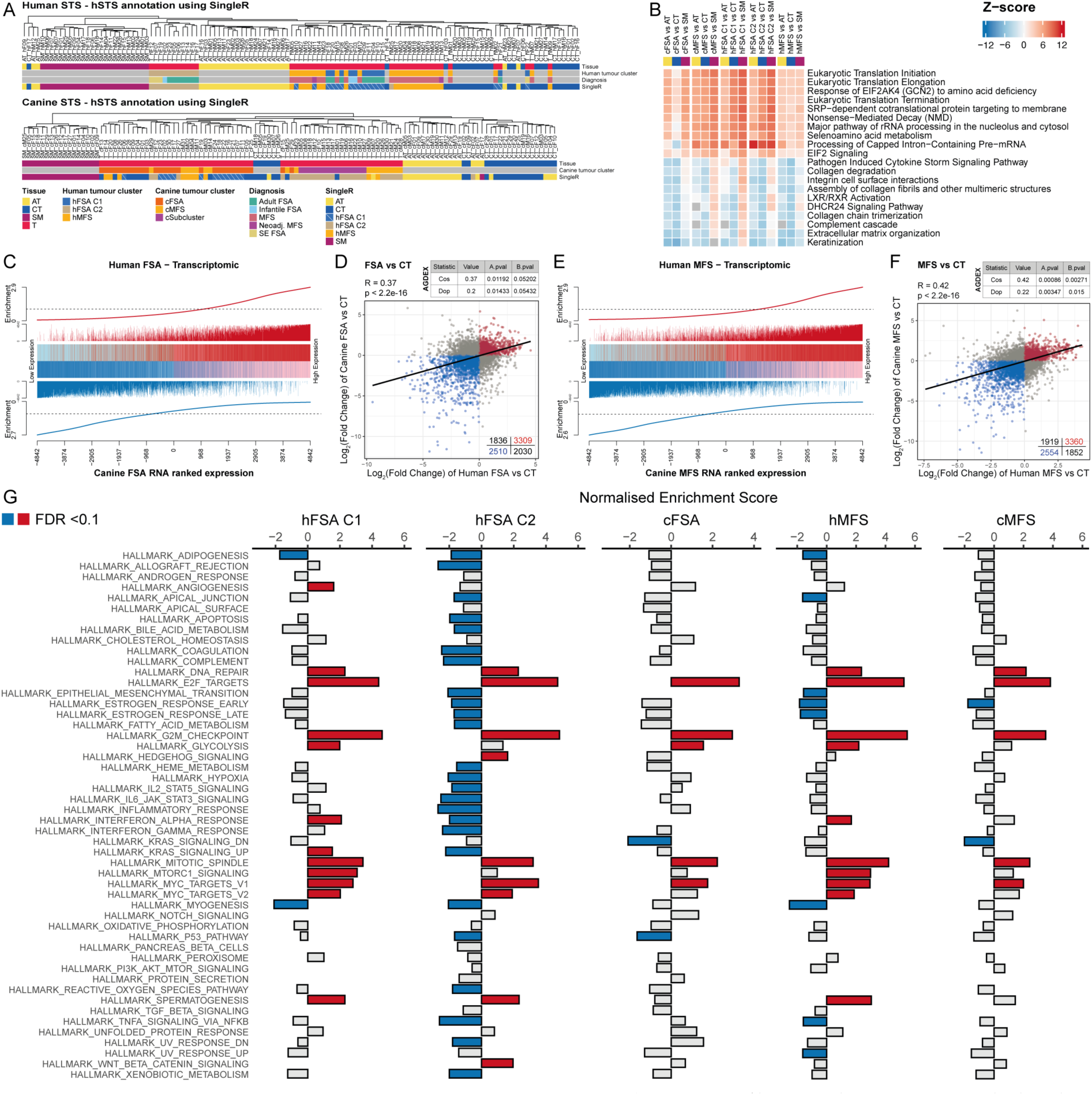
Canine and human FSA and MFS share high molecular similarity. A) Annotation of human and canine tissue samples based on the gene signature of human tissue samples using SingleR. B) Ingenuity pathway analysis displaying top10 of activated and deactivated canonical pathways using only significantly dysregulated proteins (|log2(FC)| >1 & FDR <0.05) in the comparison of tumour versus NT (AT, CT and SM). Canonical pathways ranked by the mean value in tumour versus CT. C+E) Competitive gene set testing comparing canine and human C) FSA and E) MFS. GSEA-like running sum statistic depicting the location of human genes on a ranked list of genes in the canine datasets (without cSubcluster samples). Upregulated genes are indicated as red vertical bars and downregulated genes are indicated as blue vertical bars. D+F) Correlation analysis using agreement of differential analysis (AGDEX) to compare dysregulated genes in tumour versus CT to evaluate cross-species homology in canine and human D) FSA and F) MFS (without cSubcluster samples). Depicted are the numbers of the genes in each quadrant where red dots are commonly upregulated, blue dots commonly downregulated and grey dots discordantly regulated genes. Cos, dop, Pearson correlation coefficient (R) and significance values are displayed. G) Gene set enrichment analysis of differentially expressed genes in tumour versus CT using HALLMARK gene sets. Gene sets with significant enrichment (FDR <0.1) are coloured in red if positively or blue if negatively enriched. AT = adipose tissue; cos = cosine of the angle; CT = connective tissue; dop = difference of proportions; SM = skeletal muscle.

To unravel upstream regulators influencing components of different tumour clusters, we performed transcriptional regulatory network analysis. After construction of a regulatory transcriptional network (RTN), we identified 36 and 104 regulons in the human and canine cohort, respectively (*Suppl. Fig. 15*). Unsupervised hierarchical clustering highlighted different activation of regulons and showed similar separation of tumours as ssGSEA using HALLMARK gene sets, strongly separating tumours by immune activation (*Suppl. Fig. 14A+B*). Especially the novel cSubcluster, could be strongly separated by regulon activity (*Suppl. Fig. 15B*). Comparing the activation status of the regulons between tumour and CT highlighted changed activation status (*Suppl. Fig. 15C*), including 8 regulons identified as significantly changed in both canine and human tumour clusters, with MYBL2 and FOXM1 as commonly activated (particularly involved in cell cycle progression and oncogenic processes) and IKZF1 as commonly deactivated in human and canine STS across all tumour clusters (*Suppl. Fig. 15D*). Interestingly, the higher activation status of NFIA, PRR12, TEAD2, SALL2 and ZNF362 in hFSA C2 compared to C1 highlights important transcription factors involved in stem cell maintenance, cellular differentiation and nervous system development. Contrary, the transcription factors TFEC, PRDM1, DRAP1, SPI1 and IRF8 uniquely upregulated in hFSA C1, share the regulation of immune cell function and development.

As such, these data clearly identify wide-ranging homology in gene- and protein expression patterns between canine and human MFS and FSA, leveraging dogs with spontaneous FSA and MFS as a suitable model to study the human disease.

### Cross-species molecular profiling identifies shared therapeutic targets

Both hFSA and hMFS represent tumours of fibroblastic origin. Whereas hMFS can be diagnosed based on its distinct histomorphologic features, hFSA remains a diagnosis of exclusion, lacking well-defined markers. Given the urgent need for diagnostic tools and cancer-specific therapeutic targets to improve treatment efficacy while minimizing off-target toxicity, we systematically compared transcriptional and proteomic profiles of tumour samples against matched NT (AT, CT, SM; *Suppl. Fig. 16–18*). Genes and proteins significantly upregulated (log2FC > 1, adj. p/FDR < 0.05), as well as proteins exclusively detected in tumour samples by LC–MS/MS, were considered candidate targets.

In the human tumours, transcriptomic analysis identified 1’138 differentially expressed genes (DEGs) in hFSA and 991 DEGs in hMFS (log2FC > 1 or < –1, adj. p < 0.05; *Suppl. Fig. 16A+ 17A).* At the proteomic level, 235 and 326 differentially expressed proteins (DEPs) were detected in hFSA and hMFS, respectively (log2FC > 1 or < –1, FDR < 0.05; *Suppl. Fig. 16B+17B*). Importantly, tumour-exclusive proteins were identified: 407 in hFSA (20 present in ≥80% of tumours; *Suppl. Fig. 16H*) and 480 in hMFS (53 present in ≥80% of tumours; Suppl. *Fig. 17D*), representing promising diagnostic and therapeutic candidates.

In canine tumours, 753 DEGs were identified in cFSA, 622 in cMFS, and 1’552 in the cSubcluster (*Suppl. Fig. 18A+C+E*). Proteomic analysis revealed 501, 356, and 377 DEPs in cFSA, cMFS, and cSubcluster, respectively (*Suppl. Fig. 18B+D+F*). Tumour-exclusive proteins were likewise abundant: 402 in cFSA (19 in ≥80% of tumours; *Suppl. Fig. 18G*), 566 in cMFS (84 in ≥80% of tumours, 17 in all tumours; *Suppl. Fig. 18H*), and 416 in cSubcluster (76 in ≥80% of tumours, 12 in all tumours; *Suppl. Fig. 18I*). *OSBPL6* was the only shared tumour-exclusive protein between cFSA and cMFS, while 27 proteins were common to cMFS and cSubcluster. No overlap was detected between cFSA and cSubcluster.

Comparing the findings across species, RNAseq identified 390, 864, and 467 genes significantly upregulated in hFSA C1, hFSA C2, and hMFS, respectively, with strong enrichment in cell cycle, DNA repair, and senescence pathways (*Suppl. Fig. 19*). A total of 184 genes were consistently upregulated across all human tumour subtypes (*Fig. 5A*), including *HMGA2*, *FBN2*, and *SPP1*. Notably, *FBN2* was more enriched in hFSA C2, whereas SPP1 showed higher expression in hFSA C1 and hMFS. In canine tumours, 343, 308, and 715 upregulated genes were detected in cFSA, cMFS, and cSubcluster, respectively, also enriched in cell cycle and DNA repair pathways (*Suppl. Fig. 20*). Shared gene signatures included 74 targets between cFSA and cMFS (e.g., *PLEKHG4*, *ITGA10*, *GAP43*; *Fig. 5B*), and 30 across all canine subtypes (e.g., *CELSR3*, *FZD2*, *ADGRB2*; *Fig. 5B*). Comparative cross-species analysis identified 13 commonly upregulated genes in human and canine FSA/MFS, with 6 also present in the cSubcluster, including the kinases *TK1* and *TTK* (*Fig. 5C*).

**Figure 5.**
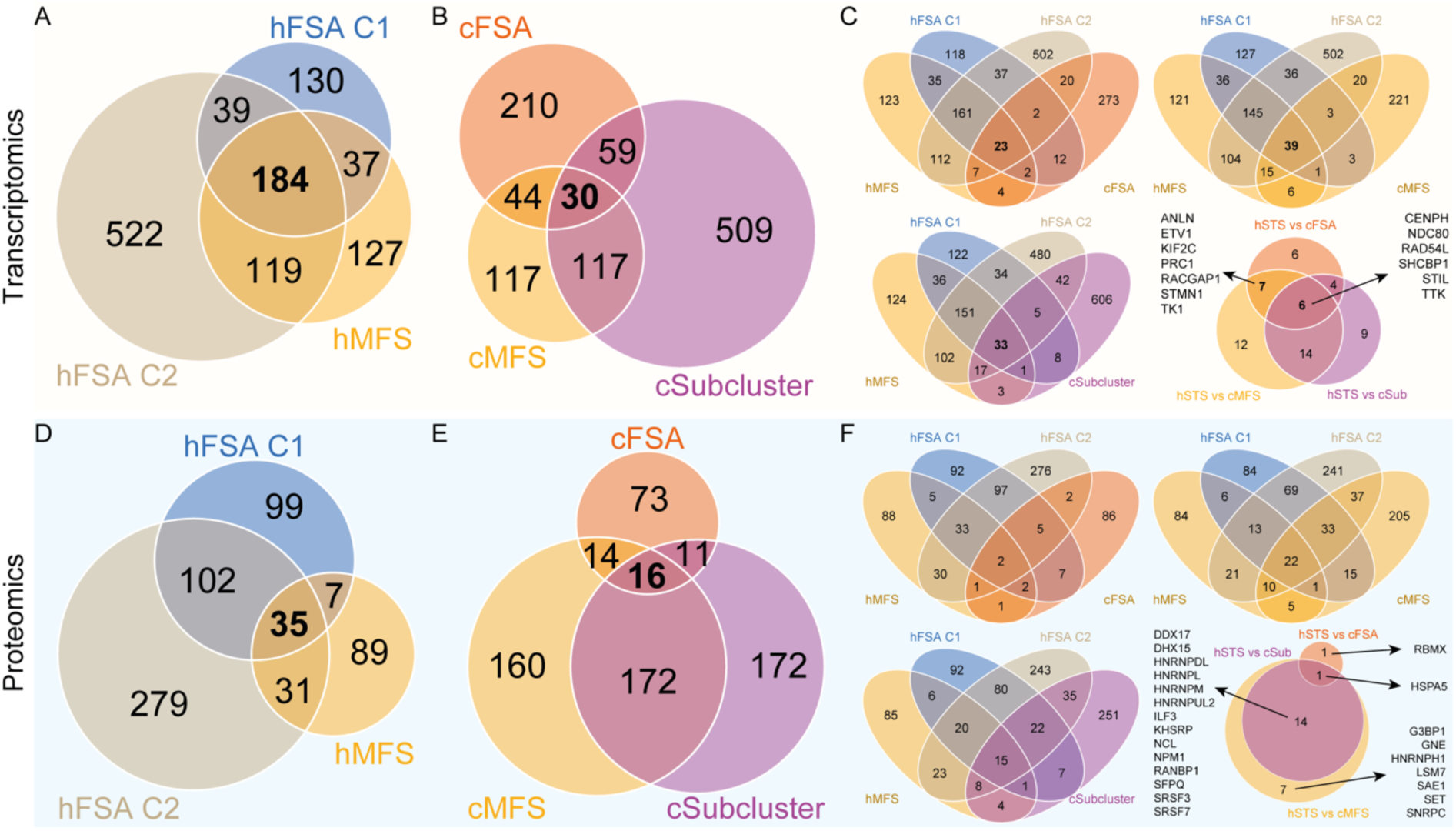
Differential gene and protein analysis reveals shared potential STS targets across species and subtypes. Venn diagram of significantly upregulated genes and proteins (log2(FC) ≥ 1 & adj. p-value/FDR <0.05) in tumour versus every NT (AT, CT or SM) to identify shared targets on A-C) transcriptomic and D-F) proteomic level. A+D) Comparison across human subtypes. B+E) Comparison across canine subtypes. C+F) Comparison across canine and human subtypes, with a final Venn diagram to identify common targets across all canine and human subtypes.

Proteomic profiling revealed 236, 397, and 109 proteins as candidates in hFSA C1, hFSA C2, and hMFS, respectively (*Suppl. Fig. 21*). Pathway enrichment highlighted MYC targets and unfolded protein response across all subtypes, with additional enrichment of immune-related programs in hFSA C1 and cell cycle pathways in hFSA C2 and hMFS. A total of 35 proteins were shared across all human tumour subtypes (e.g., *STMN1*, *LSM7*, *SAE1*; *Fig. 5D*), with 102 additional proteins shared between hFSA C1 and C2, highlighting hFSA specific targets. In canine tumours, 95, 309, and 307 candidate proteins were identified in cFSA, cMFS, and cSubcluster, respectively (*Suppl. Fig. 22*). Of these, 29 were shared across cFSA and cMFS (e.g., *COLGALT1*, *RBM3*), and 16 were shared with cSubcluster (e.g., *RBM23*, *SERPINH1*, *COLGALT1*; *Fig. 5E*).

Cross-species integration revealed *HSPA5* as a consistently upregulated protein across all human and canine tumour groups (*Fig. 5F*). Additional extracellular matrix and plasma membrane-localized proteins were identified as potential therapeutic targets, including *FSCN1*, *PLOD3*, *SPARC*, *THY1*, and *YBX1* across canine FSA and MFS. Interestingly, *FSCN1* is also upregulated in human FSA C2, hMFS, and cSubcluster, and *PLOD3* was also detected in hFSA C1, making them interesting cross-species targets for further validation. In addition, of the identified hFSA-specific targets, *PXDN* could also be identified in cFSA (Fig. 5D).

In conclusion, RNAseq and LC-MS/MS enable the identification of potential targets that could with further validation result in the development of novel therapeutic strategies. By performing molecular comparison across tissue types and species, common targets can be illuminated and tested in pre-clinical trials. This underscores canine tumours as valuable models with translational potential.

## Discussion

This multiomic portrait of FSA and MFS in human and dog aimed to characterize rare cancers in humans lacking profound molecular characterization and cross compare them to their canine counterparts. Through precise and spatial isolation of tumour and NT from FFPE blocks using LCM and subsequent analysis via RNAseq and LC-MS/MS, we have uncovered biologically/therapeutically relevant subgroups with different expression patterns and activated signalling pathways as well as susceptible targets. Our study comparing FSA and MFS between human and dogs elevates our understanding of sarcoma, leverages molecular characterization as a diagnostic tool for sarcoma and especially highlights dogs as a suitable model for rare human diseases to advance current therapeutic strategies and improve outcome in both species.

To assess whether distinct tissue types could be resolved in downstream analyses, we performed PCA and hierarchical clustering on gene and protein expression profiles (*Fig. 1, Suppl. Fig. 2+3*). Both approaches clearly separated tumour from matched NT samples, validating the effectiveness of our isolation strategy. Tumour samples clustered together irrespective of histopathological diagnosis, and tissue from different patients were highly comparable despite breed variation – a potential confounder in veterinary studies (*Suppl. Fig 1A*). Tissue-specific gene clusters further confirmed the precision of isolation, with skeletal muscle enriched for myogenesis, adipose tissue for fatty acid metabolism, and adipose/connective tissue for oestrogen response pathways (*Fig. 2+3*). In contrast, tumour samples displayed enrichment of multiple cancer-associated HALLMARK signatures, including mitotic spindle, G2M checkpoint, E2F targets, angiogenesis, and immune-related programs. Consequently, we identified hundreds of DEGs and DEPs distinguishing tumours from matched NT, providing a robust framework for the discovery of tumour-specific molecular targets (*Suppl. data 1+2*).

The incidence of human FSA has markedly declined due to refined classification of STS and recognition of genetically distinct variants, making FSA a diagnosis of exclusion ^90^. This rarity has hindered cohort assembly and comprehensive molecular characterization. Here, we present the largest hFSA cohort to date (n=23), including a primary tumour and its metastasis, 3 infantile, and 4 sclerosing epithelioid variants. Transcriptomic profiling identified two molecular clusters independent of diagnosis: an immune-enriched subgroup (hFSA C1) associated with a trend in superior disease-free survival and a neuronal-featured subgroup (hFSA C2) characterized by high proliferative activity, immune checkpoint expression and poorer survival. In addition, poor survival in the second subgroup can be linked to their gain in chromosome 12q that potentially results in upregulation of MDM2, an antagonist of the tumour suppressor gene p53 ^91^. Notably, hFSA C1 and hMFS showed strong similarities, likely reflecting shared immune activation. Major differences between hFSA and hMFS emerged in extracellular matrix composition, with hFSA showing strong upregulation of matrix metalloproteinases, highlighting potential therapeutic targets ^92^.

The canine cohorts represent the largest NGS-based, tissue-specific characterization of canine STS to date, contrasting with earlier efforts limited to bulk assessment of 16 ^93^ or 29 ^94^ mixed-subtype cases without matched NT. Prior studies therefore lacked resolution to define subtype-specific molecular signatures. By integrating tumour and matched NT, our approach enables reliable identification of tumour-specific transcriptional and proteomic alterations.

Accurate classification of STS remains challenging due to rarity, overlapping morphology, and limited pathognomonic markers ^95,96^. While human sarcoma diagnostics highlights the importance of molecular tools such as FISH and NGS – refining diagnosis in up to one-third of cases and altering treatment decisions ^97,98^ – veterinary practice still relies largely on histopathology and immunohistochemistry. Importantly, in our cohort, one cFSA and nearly half of cMFS cases formed a novel molecular different subcluster defined by the MNT–NCOA2 fusion, which may serve both as a diagnostic marker and therapeutic target in precision oncology. This underscores the value of molecular profiling in STS diagnostics, with direct implications for classification and treatment strategies. Beyond the fusion-driven cSubcluster, molecular profiling revealed a continuum between canine FSA and MFS (*Fig. 3*), analogous to the spectrum observed between hFSA C1 and hMFS. This resembles the recognized continuum between MFS and UPS/USTS ^4,13,22^. While both MFS and UPS are genomically complex, GISTIC analysis showed that hFSA C1 harbours relatively few copy number alterations. We therefore propose that hFSA C1 may represent an early stage along this continuum, with progressive genomic instability driving transition from FSA to MFS and ultimately to UPS.

The poor prognosis of STS underscores the urgent need for predictive biomarkers to stratify patients and guide biomarker-driven therapies. In our study, ssGSEA of HALLMARK gene sets consistently revealed enrichment of MYC targets V1, mitotic spindle, and protein secretion across all tumours and both species, suggesting a conserved biological core in FSA and MFS (*Fig. 2+3*). While clinical and histological features such as sex, surgical margins, anatomical site, or grade showed no association with expression patterns, ssGSEA identified four clusters defined primarily by immune- and cell cycle–related gene sets. Immune enrichment formed a continuous gradient, from absent to highly enriched tumours. Deconvolution confirmed these trends, and in human FSA/MFS high immune enrichment correlated with a trend toward longer disease-free survival, suggesting a potential protective role of immune activation. The prognostic impact is likely determined by the type and state of infiltrating immune cells. Macrophages were consistently predicted as the dominant population, in line with prior reports ^99^. Given their functional dichotomy – anti-tumorigenic M1 versus pro-tumorigenic M2 – tumour-associated macrophages (TAMs) represent an attractive therapeutic target, whether by depletion, reprogramming, or recruitment blockade ^99–101^. Conversely, dendritic cells and B cells, which were enriched only in immune-active tumours, are associated with favourable prognosis ^22,102–104^. These findings suggest that immune-active tumours may benefit from immunotherapies, such as ICI in B cell–rich sarcomas ^102^ or high expression of immune checkpoint molecules, whereas immune-desert tumours might first require sensitization with immune-modulating strategies (e.g., radiotherapy ^24^).

In canine tumours, predicted immune infiltration was generally low (*Suppl. Fig. 8),* though interpretation is limited by reliance on human-derived immune reference signatures. Since no validated canine immune reference signature exist, deconvolution may yield misleading results. Developing species-specific references is therefore critical to enable accurate prediction, immunogenomic analyses, and ultimately enhanced translational relevance of canine studies.

Although sarcomas are traditionally regarded as poorly immunogenic, subsets harbouring CNAs or mutations can trigger antitumor immunity, often counterbalanced by immune suppression ^105,106^. In hFSA C2, several CNAs were detected, but immune enrichment was absent, consistent with predicted immune checkpoint activation (*Fig. 2, Suppl. Fig. 10*). Interestingly, AHNAK (18%) and MACF1 (12%) were among the top 10 mutated genes in our human STS, which were found to be also mutated in 14 and 10% cases of canine STS ^94,107^, respectively, highlighting again cross-species STS commonalities. Of note, mutation calls from RNAseq require caution: T>C substitutions, the predominant event in our data, are often artifacts of RNA editing ^108^, while C>T transitions can result from FFPE processing ^109^. More robust analytical tools will be required to distinguish true mutations from technical artifacts ^108^.

Beyond immune heterogeneity, proliferation emerged as a key dichotomy. Enrichment of G2M checkpoint and E2F targets correlated with significantly shorter disease-free survival in human FSA and MFS (*Fig. 2*). This is consistent with Ki-67–based proliferation indices as predictors of poor outcome in other sarcomas such as dedifferentiated liposarcoma (DDLPS) ^26,110^. Conversely, low-proliferation tumours may represent less aggressive disease, raising the possibility of tailoring surgical approaches, such as narrower margins, without compromising local control. Interestingly, however, enrichment of G2M checkpoint and E2F targets did not directly correlate with tumour grade (Fig. 2G and 3F), suggesting that current assessment may not be sufficient to capture the true extent of proliferative potential in mesenchymal tumours. Although incomplete clinical data precluded similar analysis in canine tumours, however, the pattern appears suggestive. Taken together, our findings highlight immune activation and proliferative activity as one common or two independent signals associated with prognosis in FSA and MFS. While definitive conclusions are limited by retrospective design and incomplete clinical annotation in veterinary cases, similar observation have been made in human osteosarcoma patients ^56^, supporting the prognostic relevance of these pathways as promising biomarkers for clinical decision making. These results provide a rationale for prospective studies integrating molecular and immune profiling, with particular emphasis on evaluating immunotherapies such as ICI in biomarker-defined sarcoma subsets. As such, naturally occurring tumours in the context of an intact immune system underscore the utility of canine STS in comparative oncology and reinforces their translational model.

Alongside prognostic biomarkers, novel therapeutic targets are urgently needed to improve the management of STS. A critical step toward target discovery is the distinction of malignant tissue from biologically relevant NT counterparts. To this end, we incorporated skeletal muscle, adipose tissue, and connective tissue as matched controls, mirroring the anatomical context in which STS typically arise. This strategy enabled the identification of highly upregulated transcripts and proteins, as well as tumour-exclusive proteins—particularly promising candidates for precision oncology. Such tumour-restricted molecules are of special interest not only for targeted therapies but also for intraoperative near-infrared (NIR) imaging, where selective probe accumulation may facilitate real-time tumour visualization and improve surgical resection ^34,111–115^. Comparative analyses across human and canine STS highlighted several recurrent gene- and protein-level candidates (*Fig. 5*). Notably, multiple tumour-unique proteins were detected in >80% of tumours but absent from all NT, underscoring their potential as robust and specific imaging or therapeutic targets. Candidate targets include TTK protein kinase (TTK), thymidine kinase 1 (TK1), ADAM metallopeptidase with thrombospondin type 1 motif 14 (ADAMTS14), COPI coat complex subunit zeta 1 (COPZ1), protein regulator of cytokinesis 1 (PRC1), and heat shock protein family A member 5 (HSPA5). Of particular interest, HSPA5 was identified as a tumour-exclusive protein across all STS subtypes and both species. Through its involvement in the unfolded protein response, HSPA5 promotes survival, therapy resistance, and ferroptosis suppression ^116,117^, and several inhibitors are under investigation ^118^, also in human and canine OSA cell lines ^119,120^. Its cross-species tumour exclusivity and oncogenic role make it a highly attractive target for further validation.

The rarity of STS and the absence of suitable clinical models have hampered the evaluation of novel therapeutic strategies. Our findings demonstrate that the molecular architecture of STS is conserved across species – evident from shared gene expression patterns, commonly dysregulated pathways and transcription factors, and overlapping gene set enrichment profiles (*Fig. 4*) – underscoring the translational relevance of canine STS as preclinical models, particularly for testing targeted therapies and immunomodulatory interventions.

Historically, veterinary STS have been assessed almost exclusively by histopathology, an approach insufficient for capturing their biological complexity. Molecular characterization is crucial for improving diagnostic accuracy, identifying therapeutic vulnerabilities, and advancing precision medicine. Given the scarcity of research in both veterinary and human STS, the datasets generated here provide an essential foundation for future investigations. Despite their aggressive clinical course and substantial impact in companion animals, veterinary STS have lacked comprehensive molecular characterization. However, their spontaneously developing STS within an intact immune system, provide a unique opportunity to uncover novel targets and mechanisms of sarcomagenesis that can bridge the gap between experimental systems and clinical application. As such, our study addresses this gap by generating reference datasets for canine FSA and MFS, enabling refined molecular classification and cross-species comparisons. Ultimately, further validation of identified targets and their integration in clinical trials will finally unveil the massive potential of canine tumour patients as a model for human STS.

## Supporting information

Supplementary Data Canine Samples

Supplementary Data Human Samples

## Data and code availability

The mass spectrometry proteomics data have been deposited to the ProteomeXchange Consortium via the PRIDE ^121^ partner repository with the dataset identifier PXD073996 (cMFS), PXD074002 (hFSA), and PXD074073 (hMFS). The canine FSA raw MS data can also be found there (PXD029338) ^27^.The RNA sequencing raw data have been deposited in NCBI’s Gene Expression Omnibus ^122^ and are available under GEO Series accession number GSE318491 (cFSA), GSE318494 (cMFS, including re-sequenced samples and additional hFSA samples), GSE318495 (hFSA), and GSE319382 (hMFS). All other data supporting our findings is contained in the manuscript and in the supplementary figures and tables.

## Author contributions

M.C.N and E.M. planned, initiated and supervised the study. A.J., expert STS pathologist, selected clinical human cases, double-checked diagnosis and identified areas to be isolated by LCM. F.G., a nationally certified veterinary pathologist, selected clinical canine cases and identified areas to be isolated by LCM. D.F. and A.P. performed LCM and RNA extraction. D.F. and E.B. performed data visualization and prepared figures. D.F., A.J., A.F. and M.W. gathered clinical metadata. A.J. and F.G. performed immunohistochemistry. L.K. analysed all samples by LC-MS/MS and W.W. performed initial bioinformatic analysis of the LC-MS/MS data. D.F., E.B., A.P., A.S., A.K., G.R.B., V.R., N.F., A.F. and E.M. performed data analysis. E.M. and M.C.N. were responsible for study design, supervision, data analysis and funding. D.F and E.M. wrote the first draft of the manuscript. All authors read, contributed to, and approved the final manuscript.

## Acknowledgements

The authors thank the histology laboratory of the Institute of Veterinary Pathology, University of Zürich for slide preparation and technical assistance. Further thanks go to Dr. M. Dragomir for reviewing the manuscript and providing valuable feedback. Work in the lab of E.M. is financially supported by the Swiss National Science Foundation (Grants no. 310030_191988 and 10.002.567), a Center for Applied Biotechnology and Molecular Medicine (CABMM) Start-Up Grant, the Sassella Stiftung, the AKC Canine Health Foundation (CHF; grant No. 03017), the Albert-Heim-Stiftung (project No. 157), and the Wolfermann-Nägeli Stiftung. Work in the lab of M.C.N. is supported by the AKC Canine Health Foundation (CHF; grant No. 03019), the Gesellschaft für Kynologische Forschung, the Albert-Heim-Stiftung (project No. 146), the Forschungskredit der Universität Zürich, and the Bachofner legat.

