## Supplementary Figures for "Multiomic profiling of human and canine soft-tissue sarcomas reveals extensive molecular homology across species and identifies clinically relevant subgroups"

* shared first authors

\# shared senior authors

Supplementary figures

Supplementary Figure 1

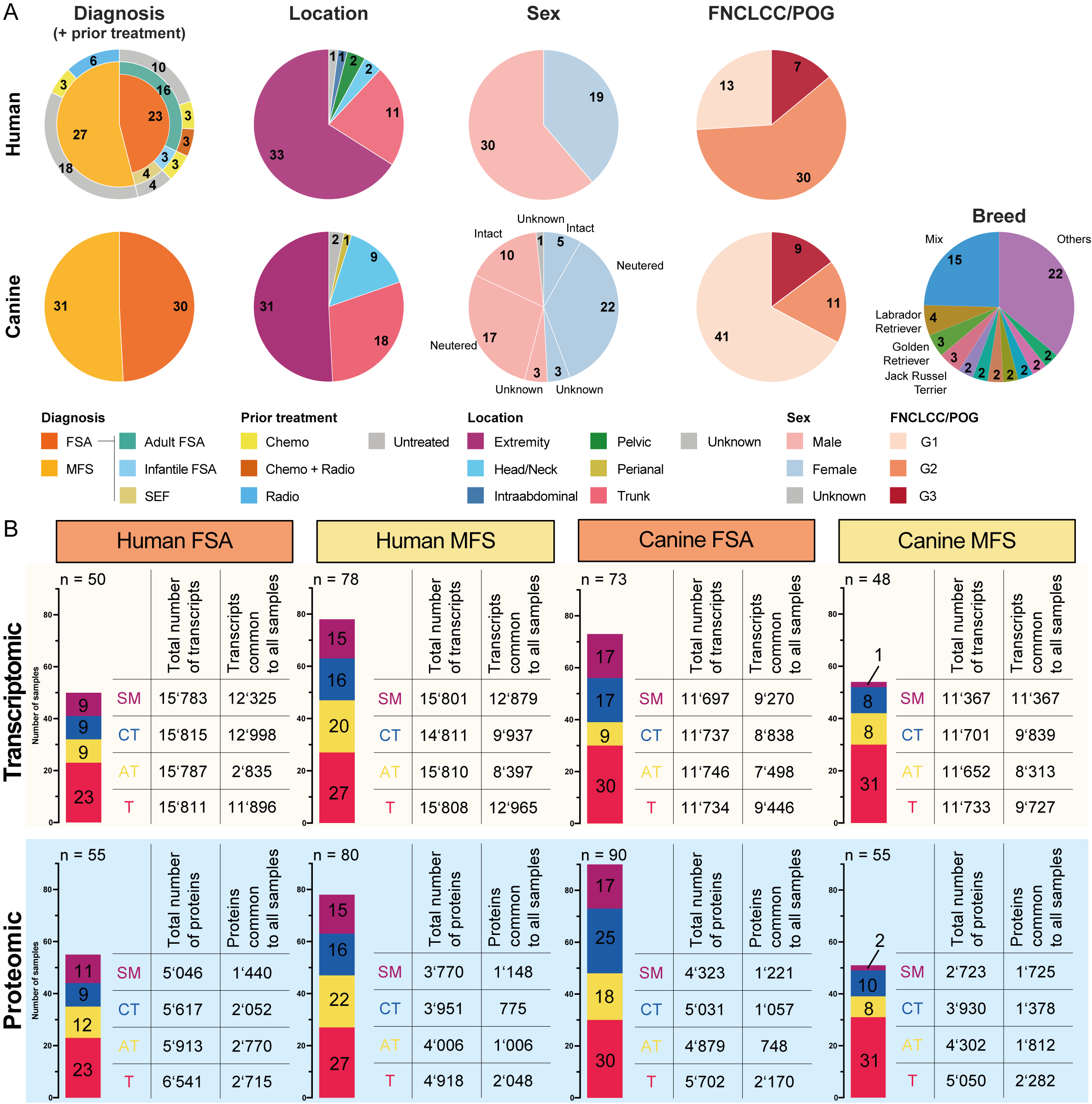

Supplementary figure 1. Overview of cases and detected transcripts and proteins.

A. Overview of clinical information for human and canine cases included in this study. SEF = sclerosing epithelioid fibrosarcoma.
B. Overview of identified transcripts and proteins per cohort and tissue type.

Supplementary Figure 2

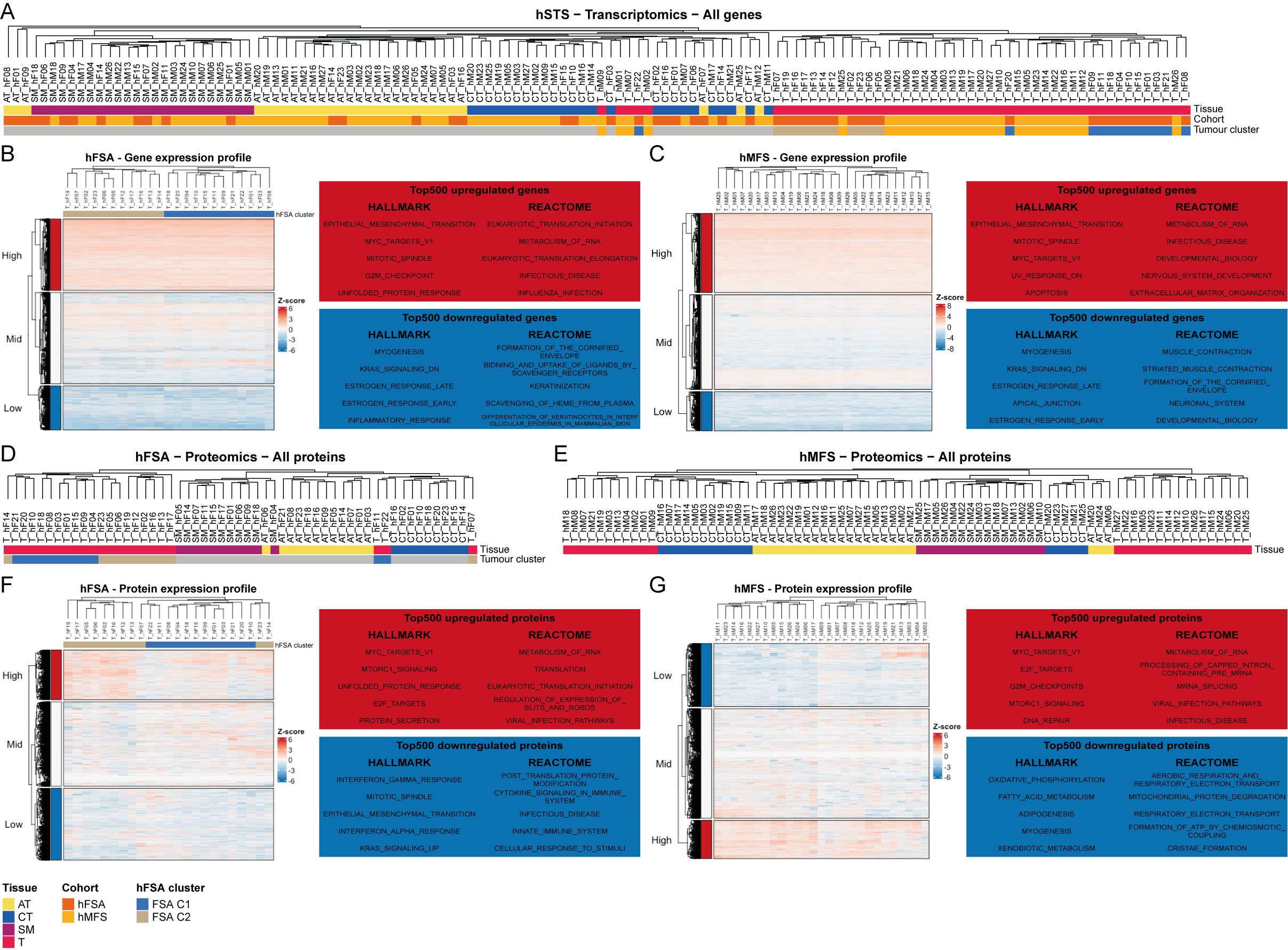

Supplementary figure 2. Hierarchical clustering of tissue samples of the transcriptomic and proteomic hSTS.

A. Hierarchical clustering of all tissue samples in the hFSA and hMFS transcriptomic cohort.

B+C.)	Heatmap of B.) hFSA and C.) hMFS cases using all genes with GSEA using HALLMARK and REACTOME pathway databases of top 500 highly and lowly expressed genes.

D+E.)	Hierarchical clustering of all tissue samples in D.) hFSA and E.) hMFS proteomic cohorts.

F+G.)	Heatmap of F.) hFSA and G.) hMFS cases using all proteins with GSEA using HALLMARK and REACTOME pathway databases of top 500 highly and lowly expressed proteins.

Supplementary Figure 3

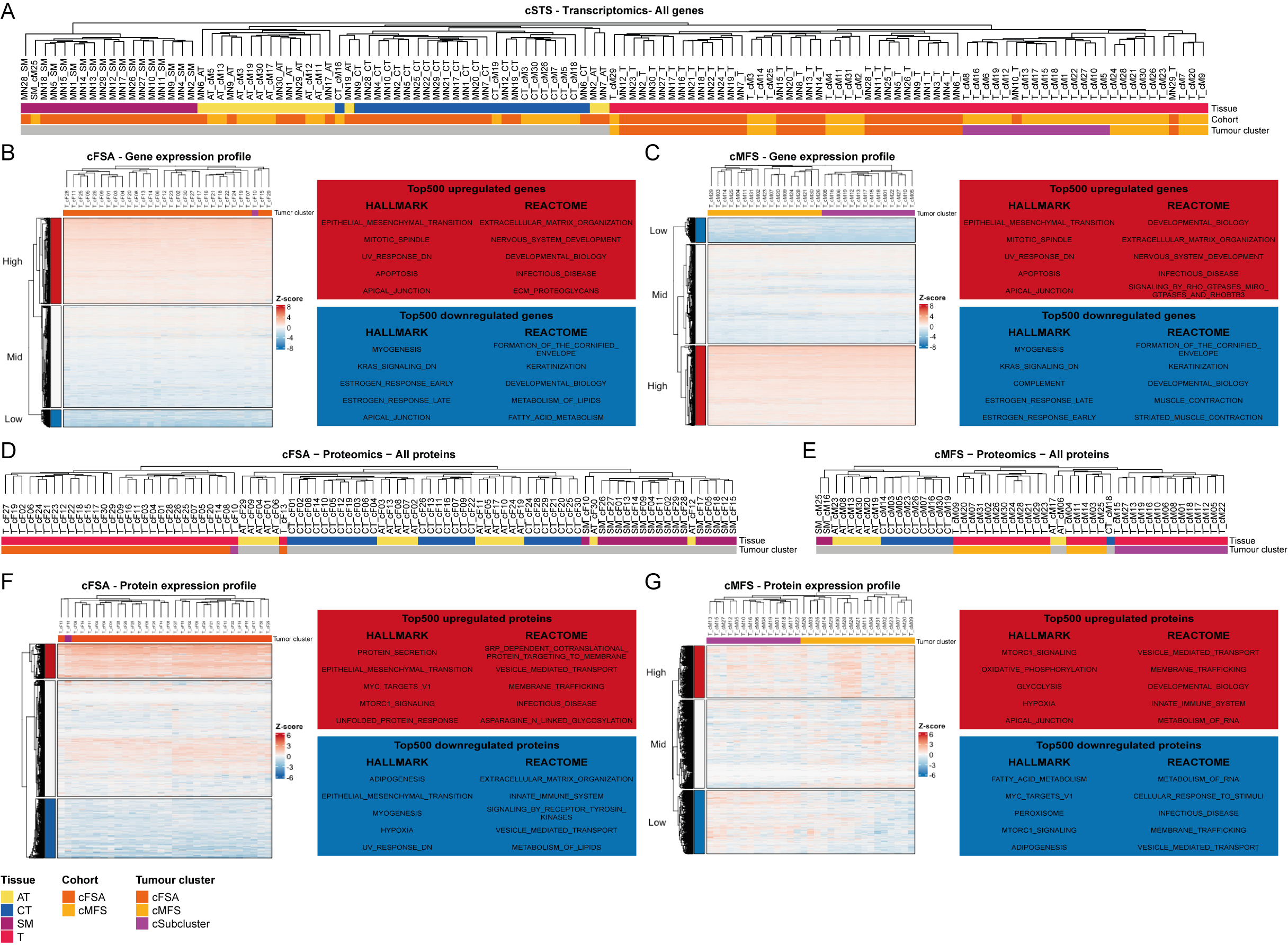

Supplementary figure 3. Hierarchical clustering of tissue samples of the transcriptomic and proteomic cSTS.

A. Hierarchical clustering of all tissue samples in the cFSA and cMFS transcriptomic cohort.

B+C.)	Heatmap of B.) cFSA and C.) cMFS cases using all genes with GSEA using HALLMARK and REACTOME pathway databases of top 500 highly and lowly expressed genes.

D+E.)	Hierarchical clustering of all tissue samples in D.) cFSA and E.) cMFS proteomic cohorts.

F+G.)	Heatmap of F.) cFSA and G.) cMFS cases using all proteins with GSEA using HALLMARK and REACTOME pathway databases of top 500 highly and lowly expressed proteins.

Supplementary figure 4

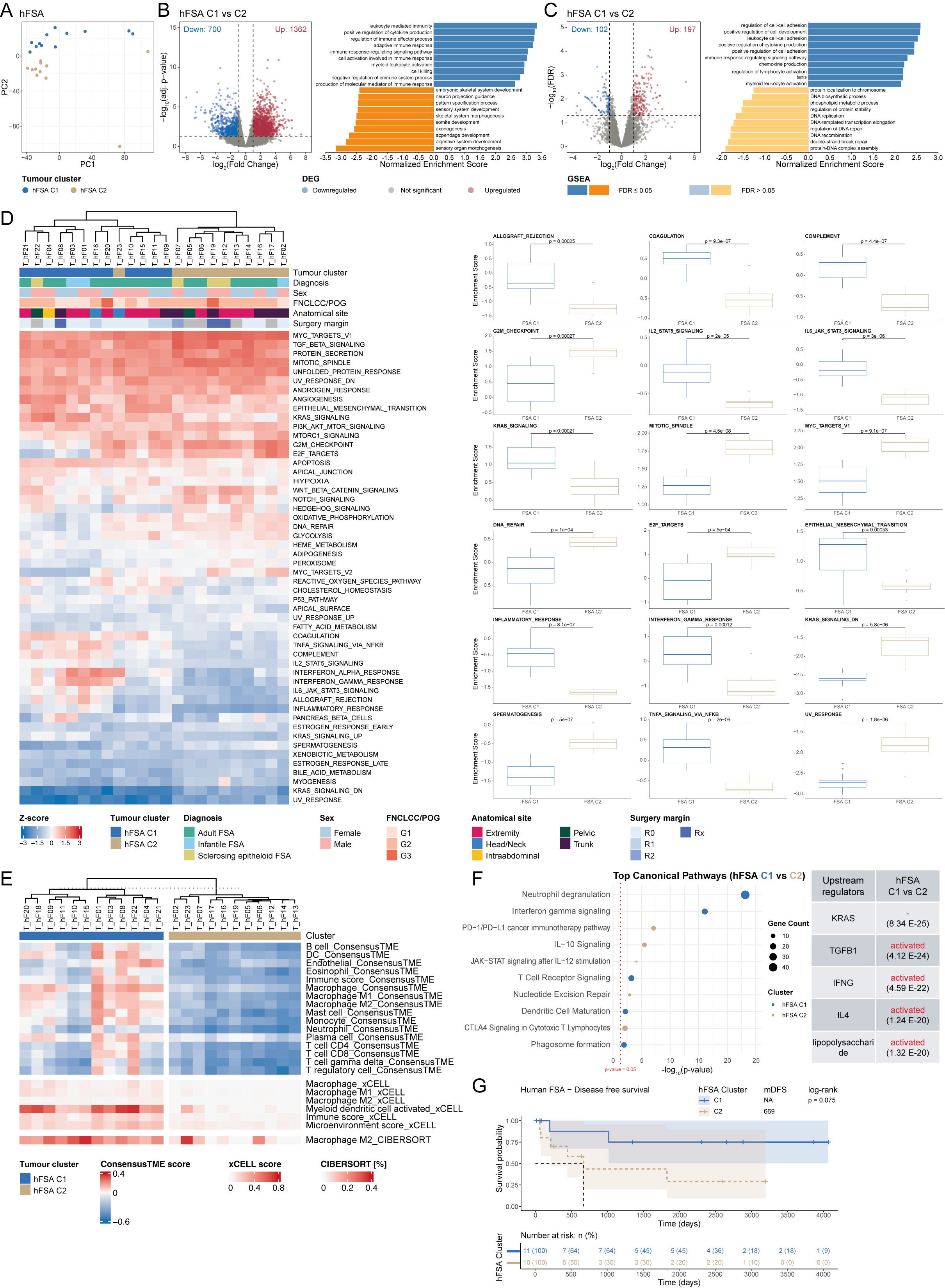

Supplementary figure 4. Molecular characterization of hFSA.

A. 2D PCA using all detected proteins in tumour from 23 human FSA.
B. Differentially expressed genes (|log_2_(FC)| ≥1 & adj. p-value <0.05) in hFSA C1 compared hFSA C2 with subsequent GSEA using Gene Ontology terms of Biological Processes.
C. Differentially expressed proteins (|log_2_(FC)| ≥1 & FDR <0.05) in hFSA C1 compared hFSA C2 with subsequent GSEA using Gene Ontology terms of Biological Processes.
D. ssGSEA using HALLMARK gene sets of all identified genes in hFSA and bar plots showing significant differentially enriched HALLMARK gene sets between hFSA C1 and C2. Student’s t-test with subsequent Bonferroni post-hoc test was performed to identify significant differentially enriched gene sets. Displayed are 18 gene sets that showed an estimated difference >0.5 or <0.5 and an adj. p-value <0.05.
E. Deconvolution analysis showing significantly different estimated immune cell populations in hFSA C1 compared to hFSA C2. Student’s t test with subsequent Bonferroni post-hoc test was performed to identify differences in estimated immune cell abundance between hFSA C1 and C2 by using an adj. p-value <0.05 as threshold.
F. IPA of canonical pathways and upstream regulators using significantly differentially expressed proteins (|log_2_(FC)| ≥1 & FDR <0.05) identified in C. Displayed are top5 most significantly canonical pathways and upstream regulators.
G. Kaplan-Meyer plot comparing disease free survival between hFSA C1 and C2, indicating a survival advantage for patients with tumours belonging to C1.

Supplementary figure 5

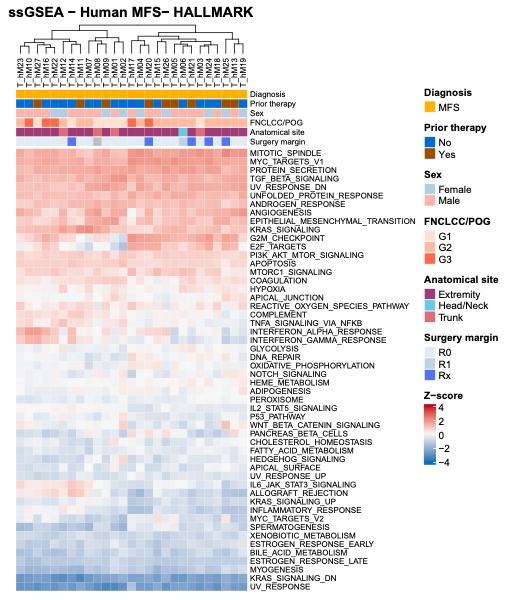

Supplementary figure 5. Molecular characterization of hMFS.

ssGSEA of hMFS cases using the HALLMARK gene sets of all identified genes reveals differences in immune- and cell cycle-related gene sets.

Supplementary figure 6

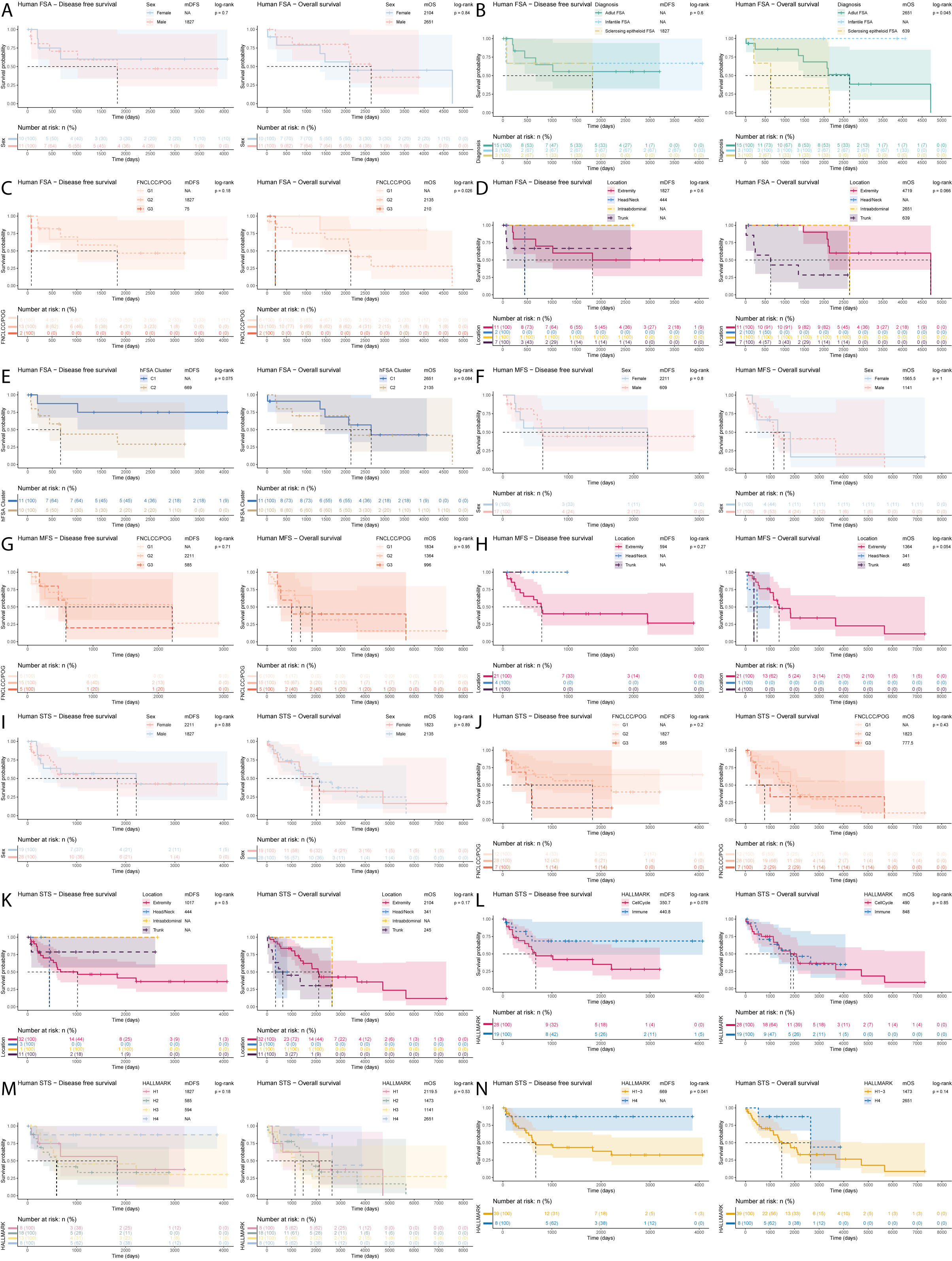

Supplementary figure 6. Kaplan Meier plots for identification of survival differences in hSTS

Kaplan Meier plots were used to identify differences in overall survival (OS) and disease-free survival (DFS) in different groups of hFSA (A-E), hMFS (F-H), or both merged hSTS (I-N) comparing A+F+I) Sex, B) Diagnosis, C+G+J) FNCLCC/POG grading, D+H+K) Tumour location, E) Cluster, L-N) HALLMARK cluster. Log-rank test was used as statistical method to evaluate difference in median DFS (mDFS) and median OS (mOS).

Supplementary figure 7

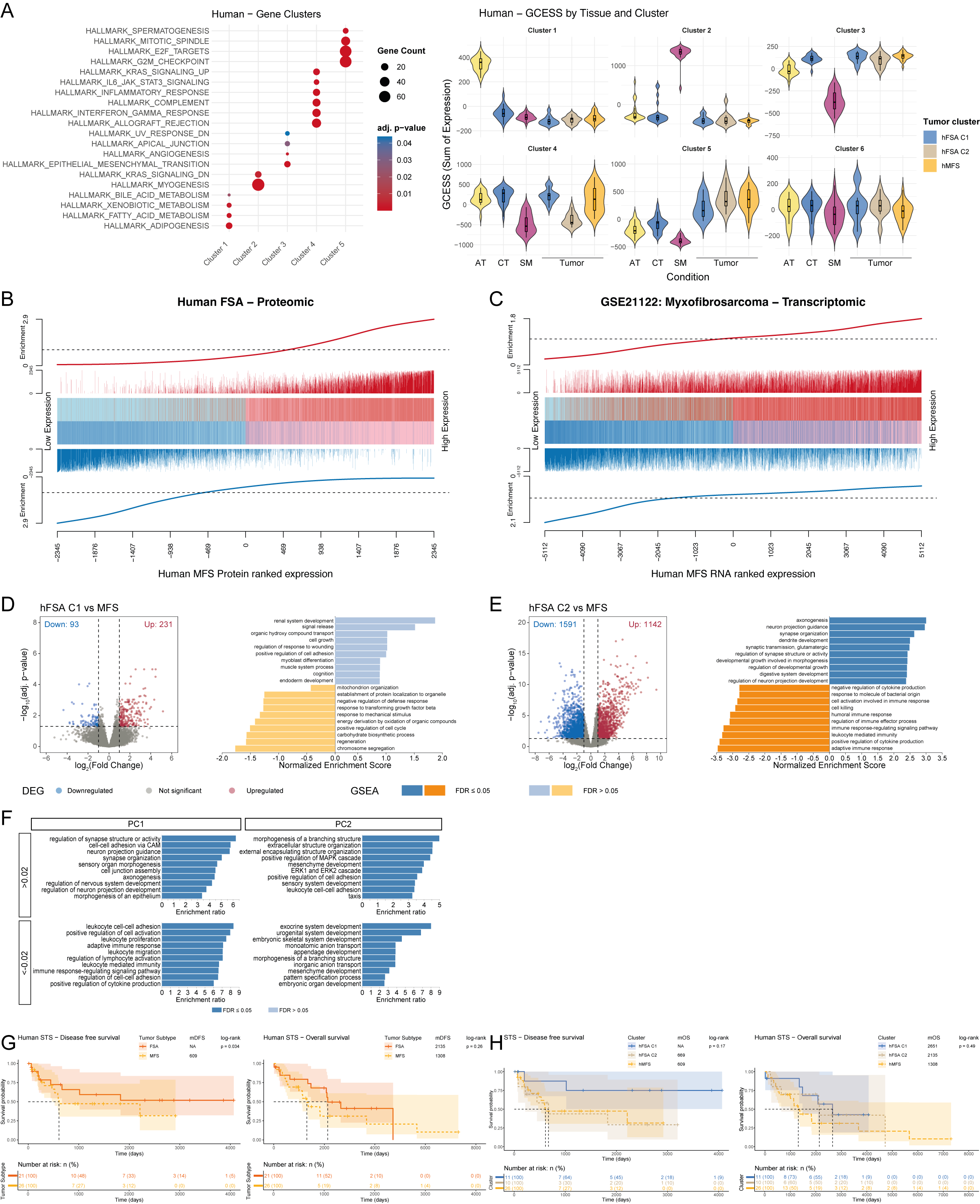

Supplementary figure 7. Molecular comparison of hFSA and hMFS.

A. GSEA of GCESS clusters identified in hSTS using HALLMARK pathways. GCESS cluster 6 did not show significant enrichment for any HALLMARK gene set and was not observed to be differently enriched between any tissue group.

B+C.	Competitive gene set testing to compare B.) our proteomic data of human MFS and human FSA and C.)

our transcriptomic hMFS data to an external transcriptomic human MFS dataset (GSE21122). GSEA-like running sum statistic depicting the location of human FSA proteins or human MFS genes (GSE21122) on a ranked list of genes in our human MFS datasets. Upregulated genes are indicated as red vertical bars, downregulated genes are indicated as blue vertical bars.

D+E.	Differentially expressed genes (|log_2_(FC)| ≥1 & adj. p-value <0.05) in D.) hFSA C1 compared to hMFS and E.) hFSA C2 compared to hMFS with subsequent GSEA using Gene Ontology terms of Biological Processes.

H. Bar plots showing the results of ORA of the first two principal components of PCA in Figure 1F using all genes with a rotation bigger than 0.02 and smaller -0.02. ORA was performed using the Gene Ontology terms of Biological Processes (no redundant). The first principal component shows clear enrichment for neural development and immune response, while the second principal component shows strong enrichment for gene sets of cell regulatory pathways and developmental systems.

G+H.	Kaplan-Meier plots comparing disease-free survival (DFS) and overall survival (OS) between G) hFSA

and hMFS and H) hFSA C1, C2 and hMFS. Log-rank test was used as statistical method to evaluate difference in median DFS (mDFS) and median OS (mOS). While hFSA cases included in our cohort seem to have a significant longer disease-free survival time than the hMFS cases, an even more pronounced better disease-free survival is observed for hFSA C1.

Supplementary Figure 8

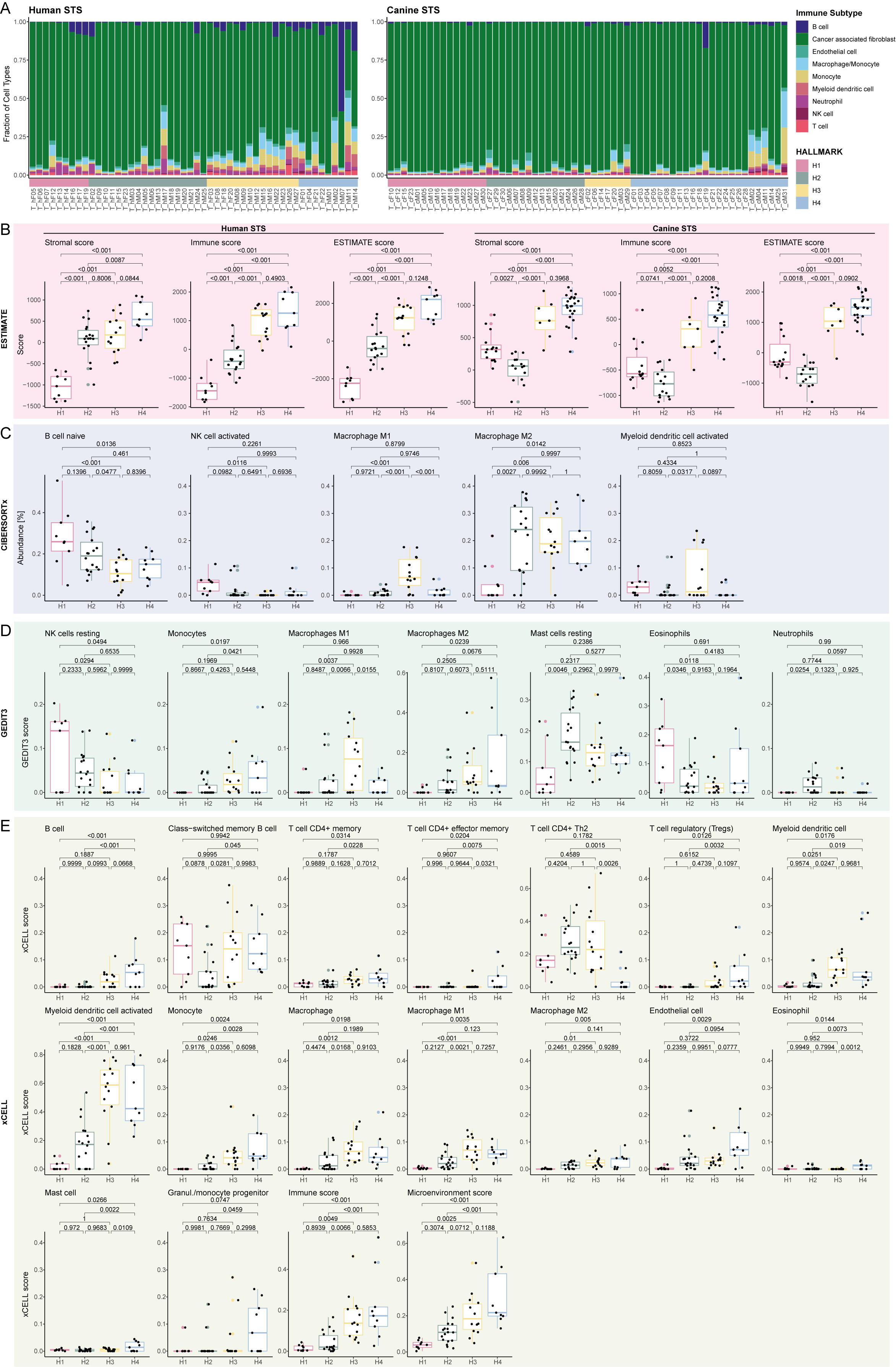

Supplementary Figure 8. Deconvolution of human and canine STS for prediction of immune cell populations.

A. Deconvolution of immune cell abundances in percentage using MCPcounter in human and canine STS tumour samples with annotation by HALLMARK clusters.
B. Deconvolution using ESTIMATE to estimate stromal, immune and ESTIMATE score in human and canine STS tumour samples.

C-E.	Deconvolution of immune cells in human STS using C) CIBERSORTx, D) GEDIT3 and E) xCell. Displayed are all immune cell subtypes with significant differences (adj. p value < 0.05) between HALLMARK clusters. For statistical analysis abundance or scores were compared using ANOVA with correction for multiple testing using post hoc Tukey test.

Supplementary Figure 9.

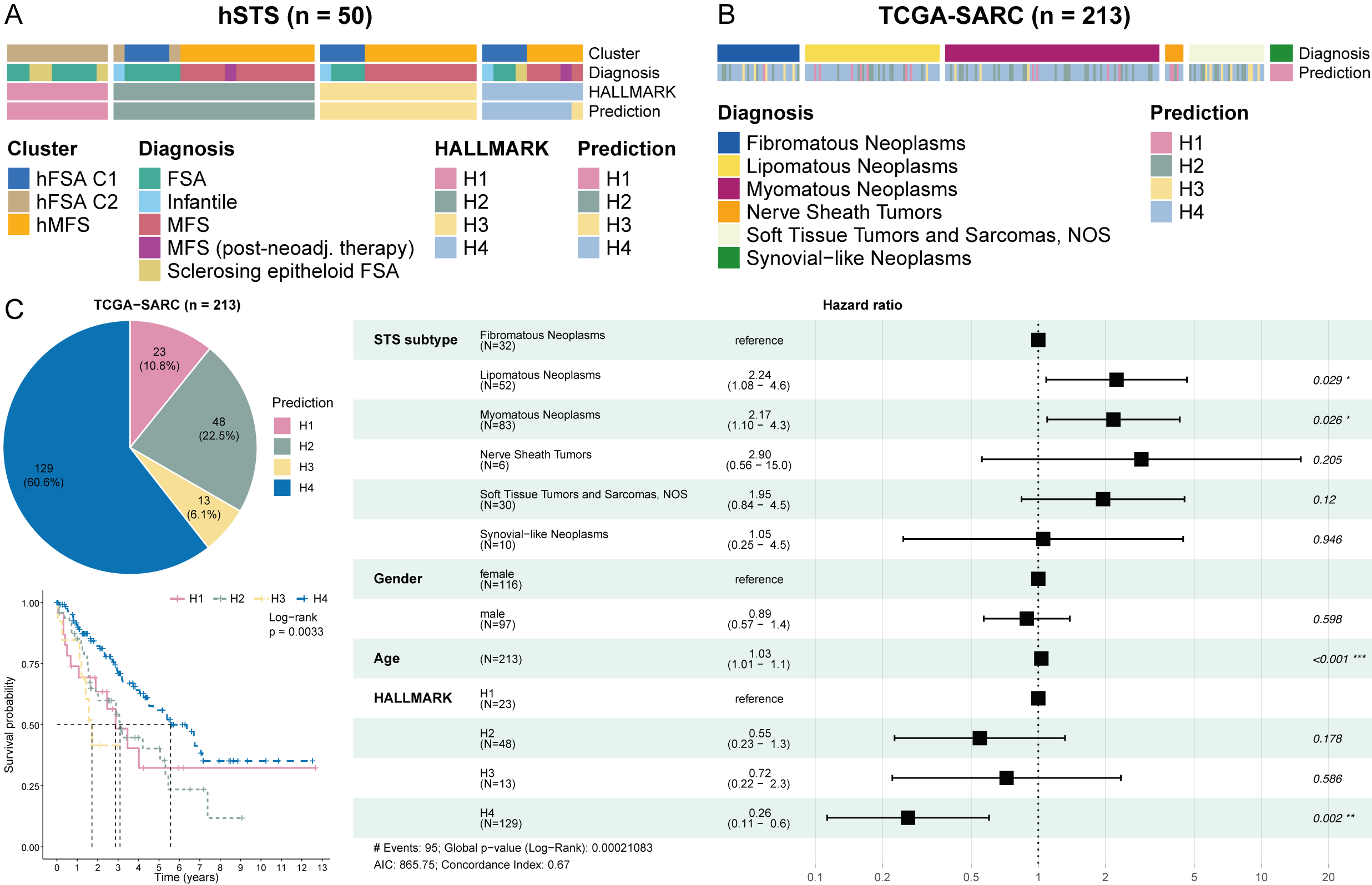

Supplementary Figure 9. External validation of prognostic value of the HALLMARK clustering.

A. Annotation of human STS tumour samples based on the gene signature of the 4 identified HALLMARK clusters using SingleR.
B. Annotation of human STS tumour samples of the TCGA-SARC data set based on the gene signature of the 4 identified HALLMARK clusters using SingleR.
C. PieChart showing the abundance of samples classified to the respective HALMARK cluster together with a Kaplan-Meier plot and a Forest plot indicating superior overall survival for tumours classified as cluster H4.

Supplementary Figure 10

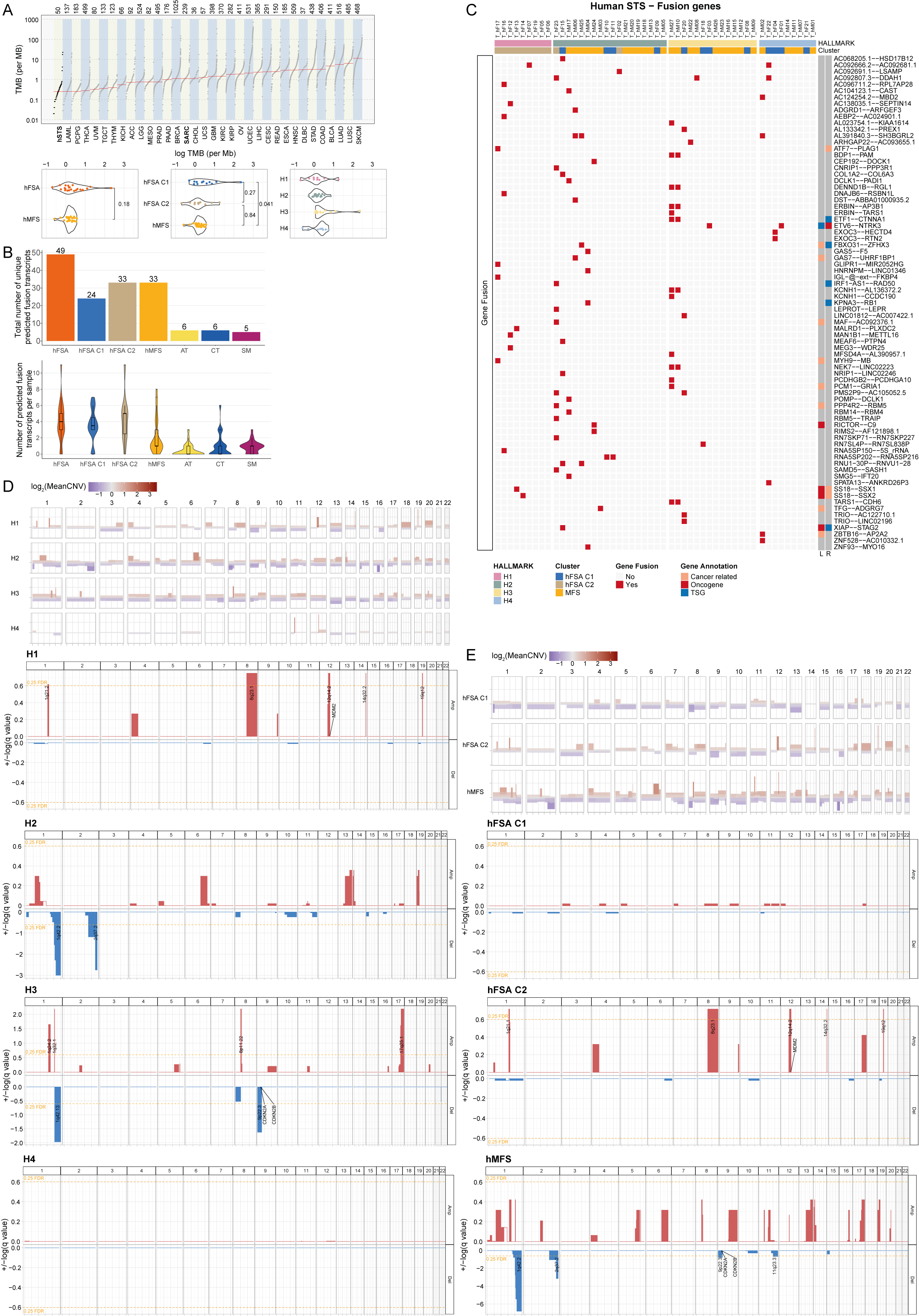

Supplementary Figure 10. Genomic alterations in human FSA and MFS predicted using RNAseq data.

A. Assessment of tumour mutational burden (TMB) across the cohort of human STS compared to TCGA cohorts and separated by different groups (Subtype, FSA Cluster and HALLMARK Cluster).
B. Number of identified gene fusions across tissues/samples.
C. All detected gene fusions in tumour tissues after removal of gene fusions also found in normal tissue.

D+E.	GISTIC analysis of CNV between D) HALLMARK clusters and E) hFSA C1, hFSA C2 and hMFS. Segments were considered as significantly amplified or deleted within a group of tumour samples using a FDR < 0.25.

TSG = tumour suppressor gene.

Supplementary Figure 11.

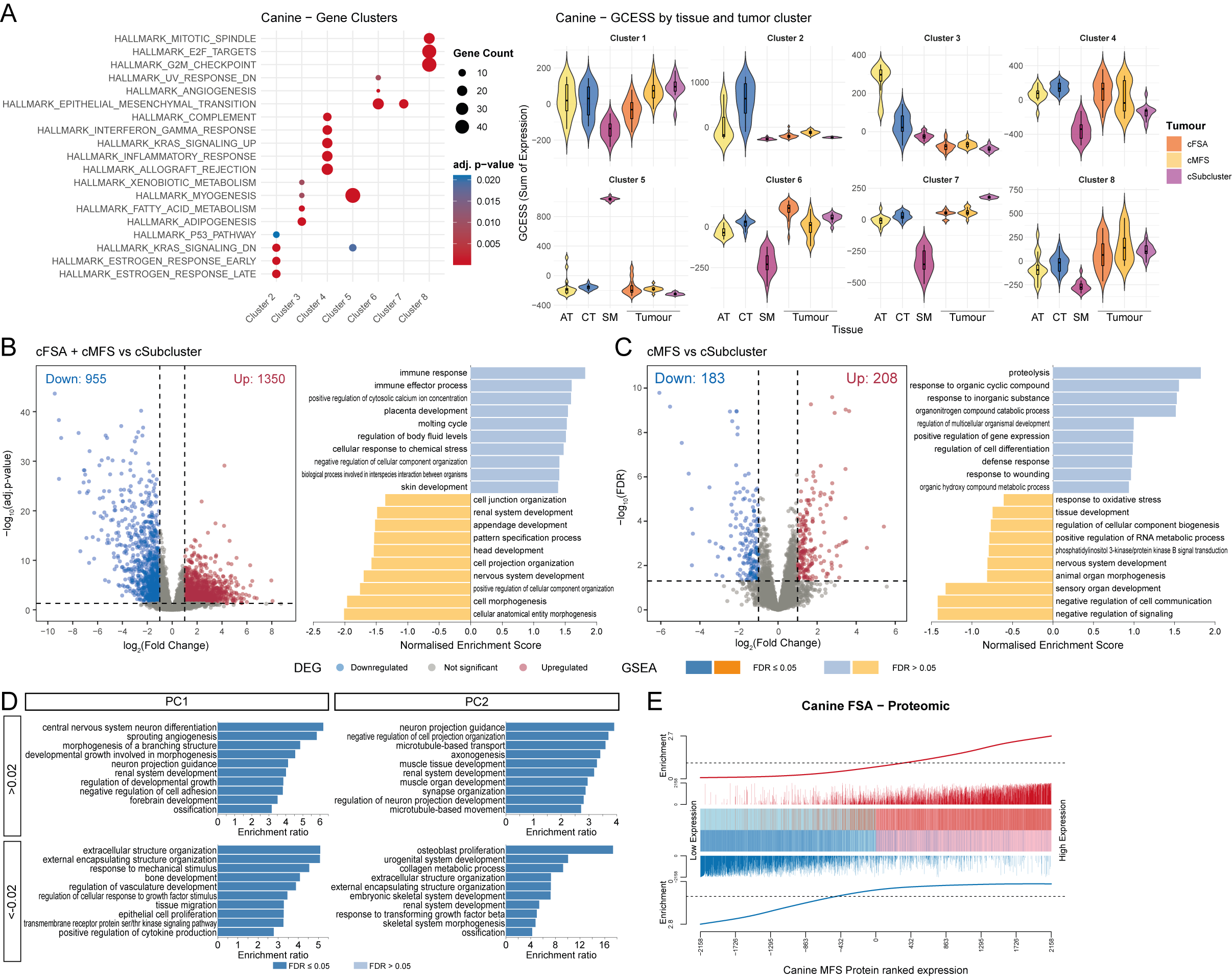

Supplementary Figure 11. Comparison of canine STS entities.

A. GSEA of gene clusters identified in Fig 3C using the HALLMARK gene sets and bar plots showing GCESS of each gene cluster by tissue type and tumour cluster. GCESS cluster 1 did not show significant enrichment for any HALLMARK gene set.
B. Differentially expressed genes (|log_2_(FC)| ≥1 & adj. p-value <0.05) in cFSA + cMFS vs cSubcluster with subsequent GSEA using Gene Ontology terms of Biological Processes.
C. Differentially expressed proteins (|log_2_(FC)| ≥1 & FDR <0.05) in cMFS vs cSubcluster with subsequent GSEA using Gene Ontology terms of Biological Processes.
D. Bar plots showing the results of ORA of the first two principal components of PCA in Figure 3B using all genes with a rotation bigger than 0.02 or smaller than -0.02. ORA was performed using the Gene Ontology terms of Biological Processes (no redundant). PC1 shows clear enrichment in nerval development and angiogenesis, while PC2 shows strong enrichment for gene sets of developmental systems.
E. Competitive gene set testing to compare cFSA and cMFS (without cSubcluster) on protein level. GSEA-like running sum statistic depicting the location of genes in one group on a ranked list of genes in the other group. Upregulated genes are indicated as red vertical bars, downregulated genes are indicated as blue vertical bars.

Supplementary Figure 12.

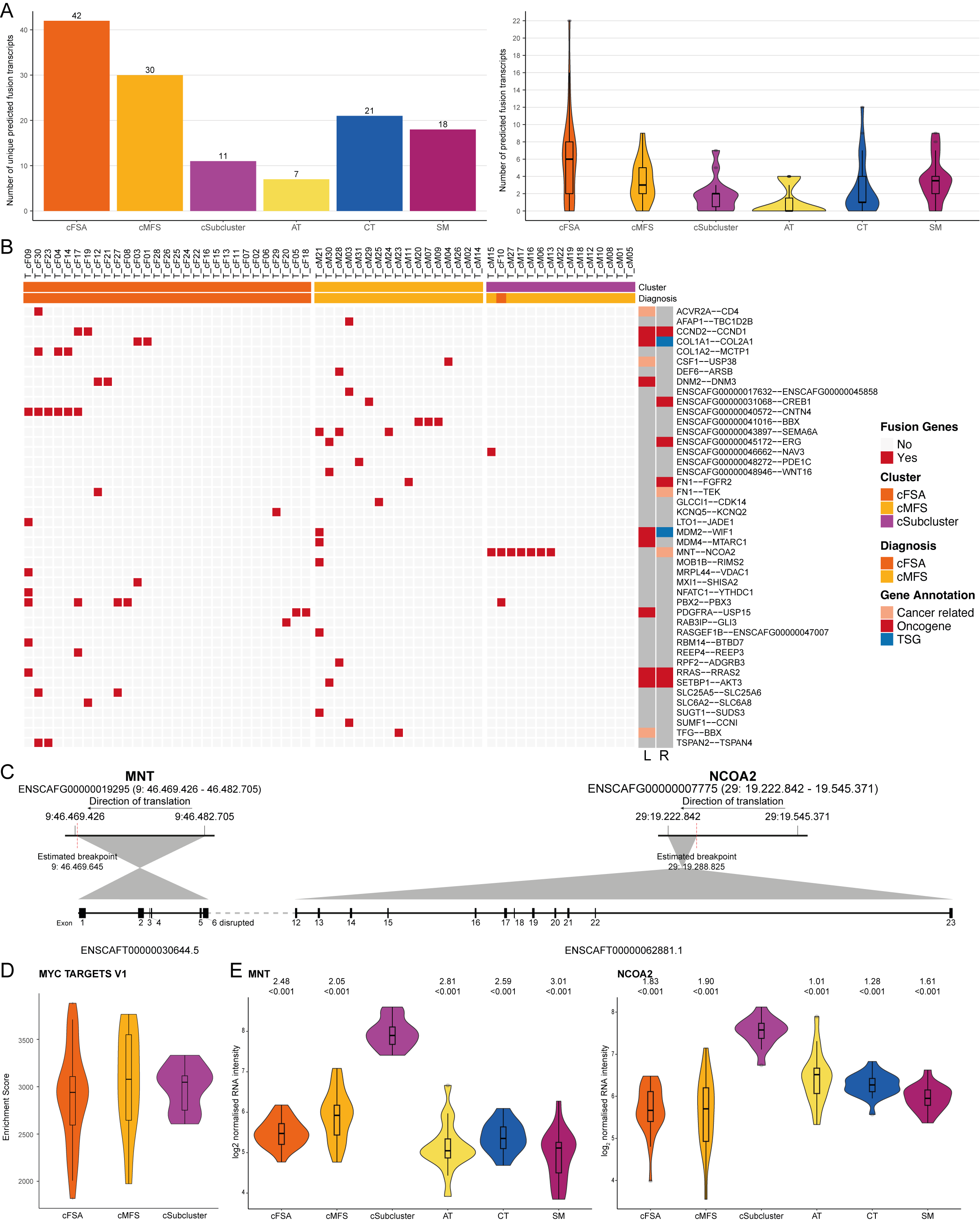

Supplementary Figure 12. Gene fusions in canine STS.

A. Overview of total number of predicted gene fusions across tissue type and per sample.
B. All detected gene fusions in canine tumour samples after removal of gene fusions also found in normal tissue.
C. Displayed are the MNT and NCOA2 gene region with indicated breakpoint and resulting transcripts with disruption of exon 6 of MNT and continuation with exon 12 in NCOA2.
D. Comparison of enrichment score in Myc Targets V1 (HALLMARK gene sets) between cFSA, cMFS and cSubcluster using single sample gene set enrichment analysis.
E. Comparison of gene expression level of MNT and NCOA2 in cFSA, cMFS and cSubcluster. Displayed values indicate log_2_FC in gene expression and adj. p-values of cSubcluster compared to cFSA, cMFS and NT (AT, CT and SM).

AD1/AD2 = autonomous transcription activation domain 1/2, BHLH = basic helix-loop-helix, CID = CBP interaction domain, NID= nuclear receptor interaction domain, PAS = Per-Arnt-Sim domain, Q rich = glutamine rich domain, TSG = tumour suppressor gene.

Supplementary Figure 13.

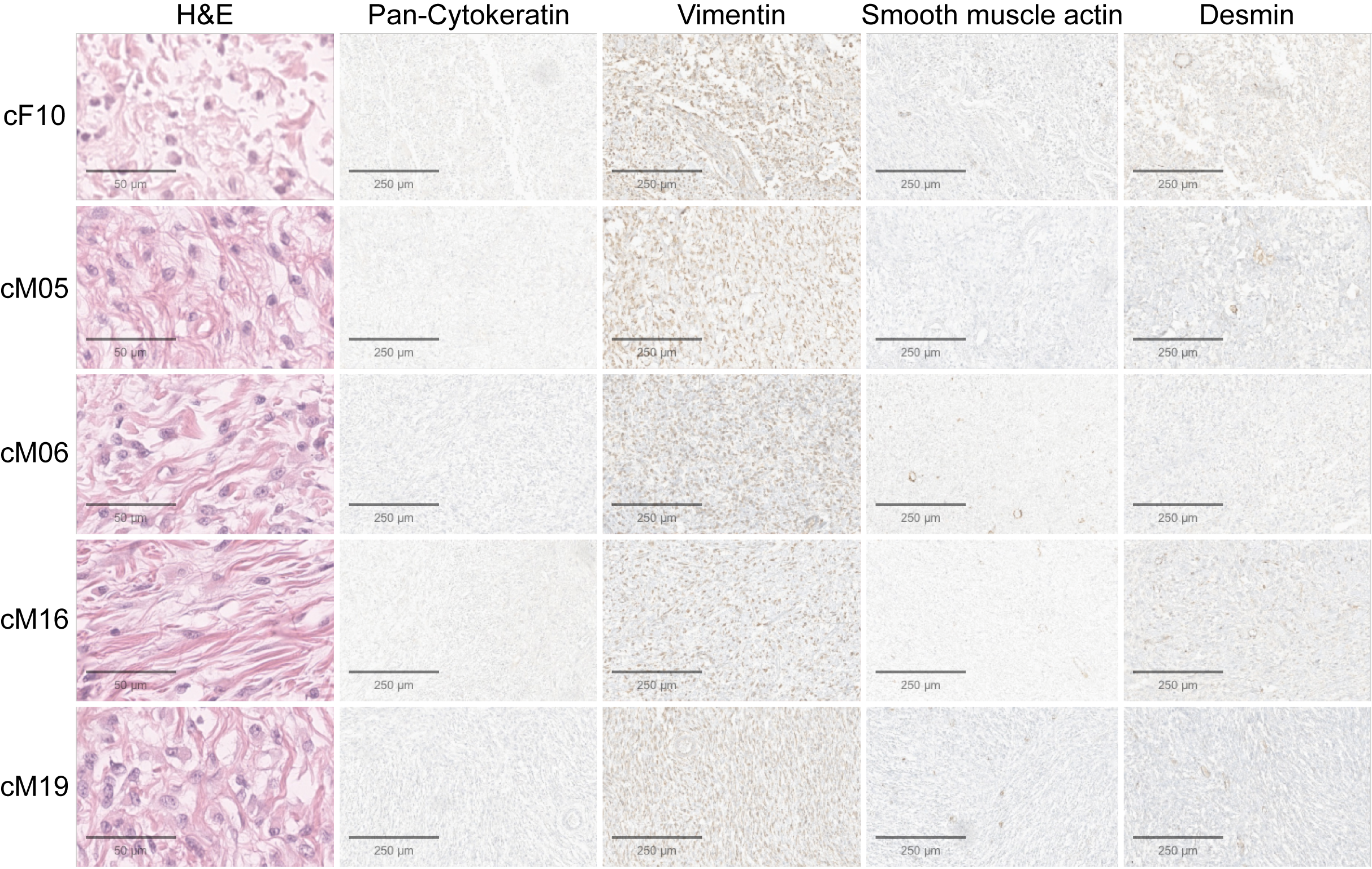

Supplementary Figure 13. Immunohistochemical evaluation of 5 cases belonging to the cSubcluster.

Cases belonging to the cSubcluster were re-evaluated based on their H&E staining together with immunohistochemistry using Pan-Cytokeratin, Vimentin, Smooth muscle actin and Desmin. Of all markers tested, all 5 cases stained only positive for Vimentin indicating a mesenchymal tumour further excluding a tumour of muscular origin.

Supplementary Fig 14.

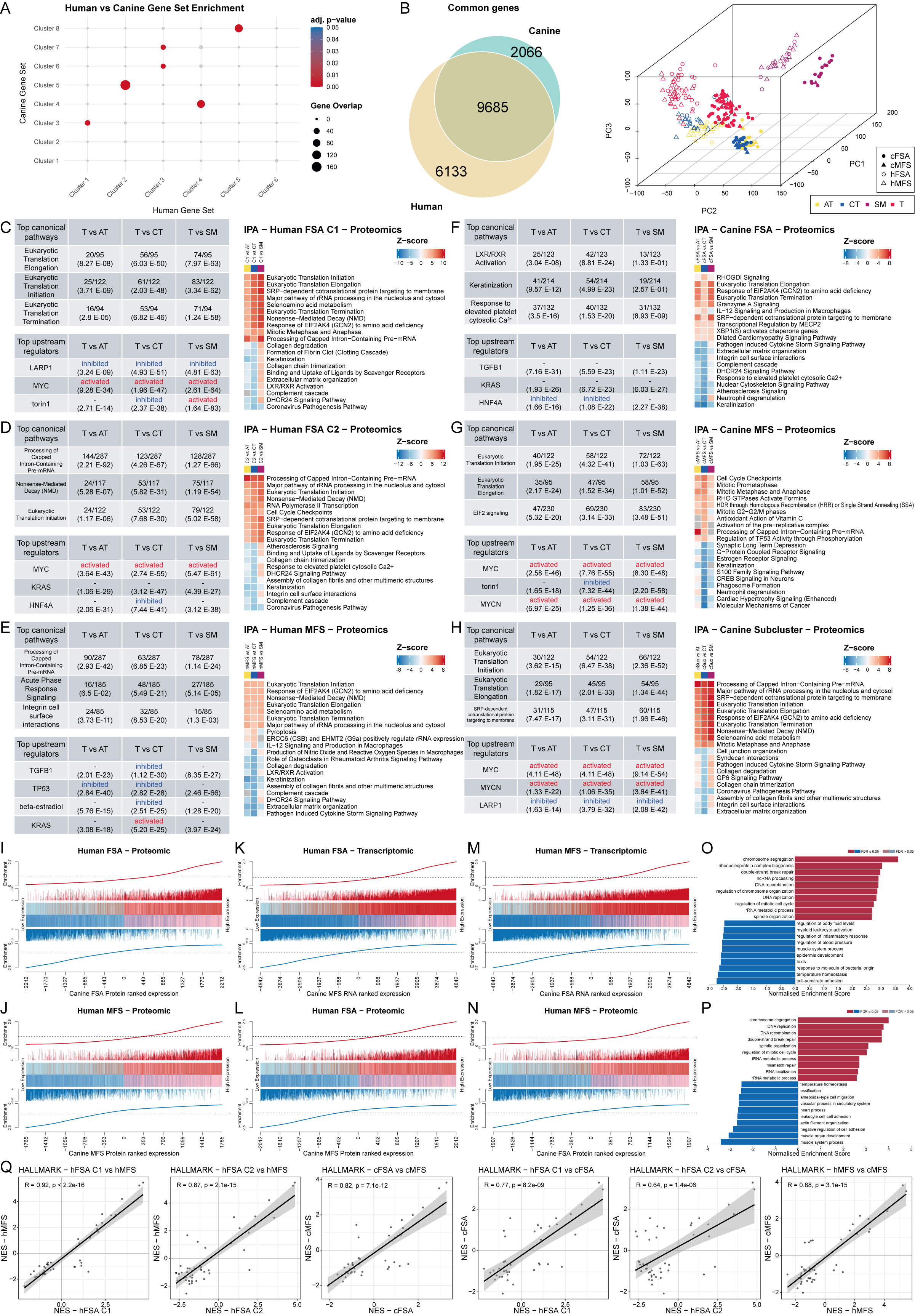

Supplementary Figure 14. Comparison between human and canine STS.

A. Pairwise overlap analysis using Fisher’s Exact Test to check enrichment for each human and canine set pair.
B. Overlap of common transcripts identified in the canine and human cohort after filtering out lowly abundant genes used for PCA showing similar tissue separation pattern in the canine and human cohort. Tissue samples are separated by tissue type, while canine and human tissue samples are separated by PC2.

C - H.	Ingenuity pathway analysis (IPA) showing top 3 canonical pathways and upstream regulators detected in T vs AT, CT and SM, as well as the activation status of top 10 positively and negatively activated canonical pathways of A) hFSA C1, B) hFSA C2, C) hMFS, D) cFSA, E) cMFS and F) cSubcluster using the proteomic data as input. Values in the table represent the number of detected targets/number of targets attributed to the respective dataset and values in the brackets represent p-values.

I - N.	Competitive gene set testing to compare A) hFSA versus cFSA and B) hMFS versus cMFS on protein level, as well as comparing hFSA versus cMFS on I) RNA and J) protein level, and hMFS versus cFSA on K) RNA and L) protein level. GSEA-like running sum statistic depicting the location of genes in one group on a ranked list of genes in the other group. Upregulated genes are indicated as red vertical bars, downregulated genes are indicated as blue vertical bars.

O + P.	GSEA using Gene Ontology terms of Biological Processes of all genes identified as commonly deregulated in AGDEX.

Q.	Correlation analysis of GSEA results using HALLMARK gene sets comparing human and canine entities.

Supplementary Figure 15.

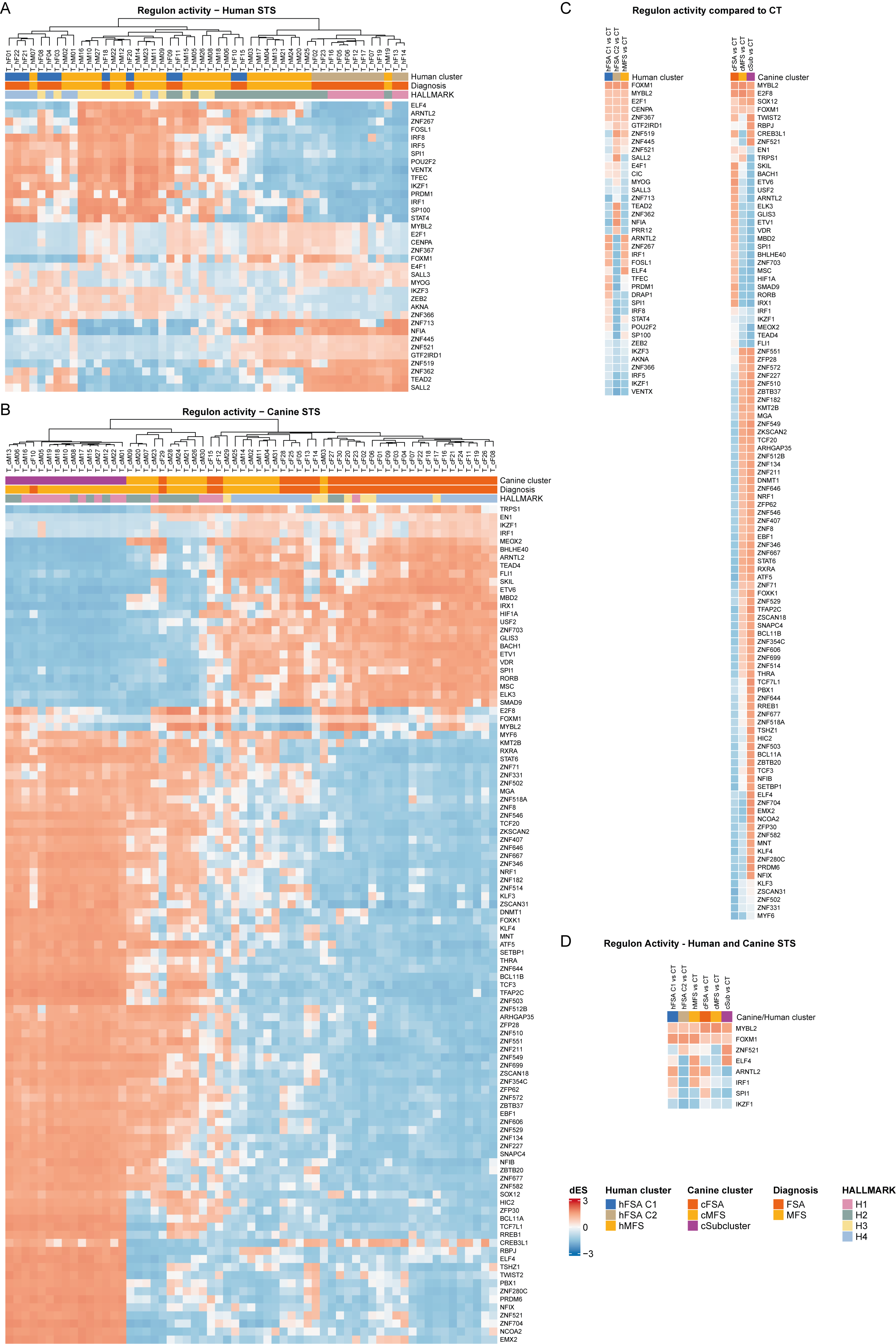

Supplementary Figure 15. Changes in regulon activity in human and canine STS.

A + B.	Regulon activity in A) human and B) canine STS using the mean estimated activation status across all tumour samples showing similar cluster pattern compared to HALLMARK clusters separating tumours in immune active and proliferative cases.

C.	Regulon activity of human and canine STS tumour cases compared to CT.

D.	Commonly identified regulons between human and canine STS tumour samples. Activation level is displayed in comparison to CT.

Supplementary Figure 16.

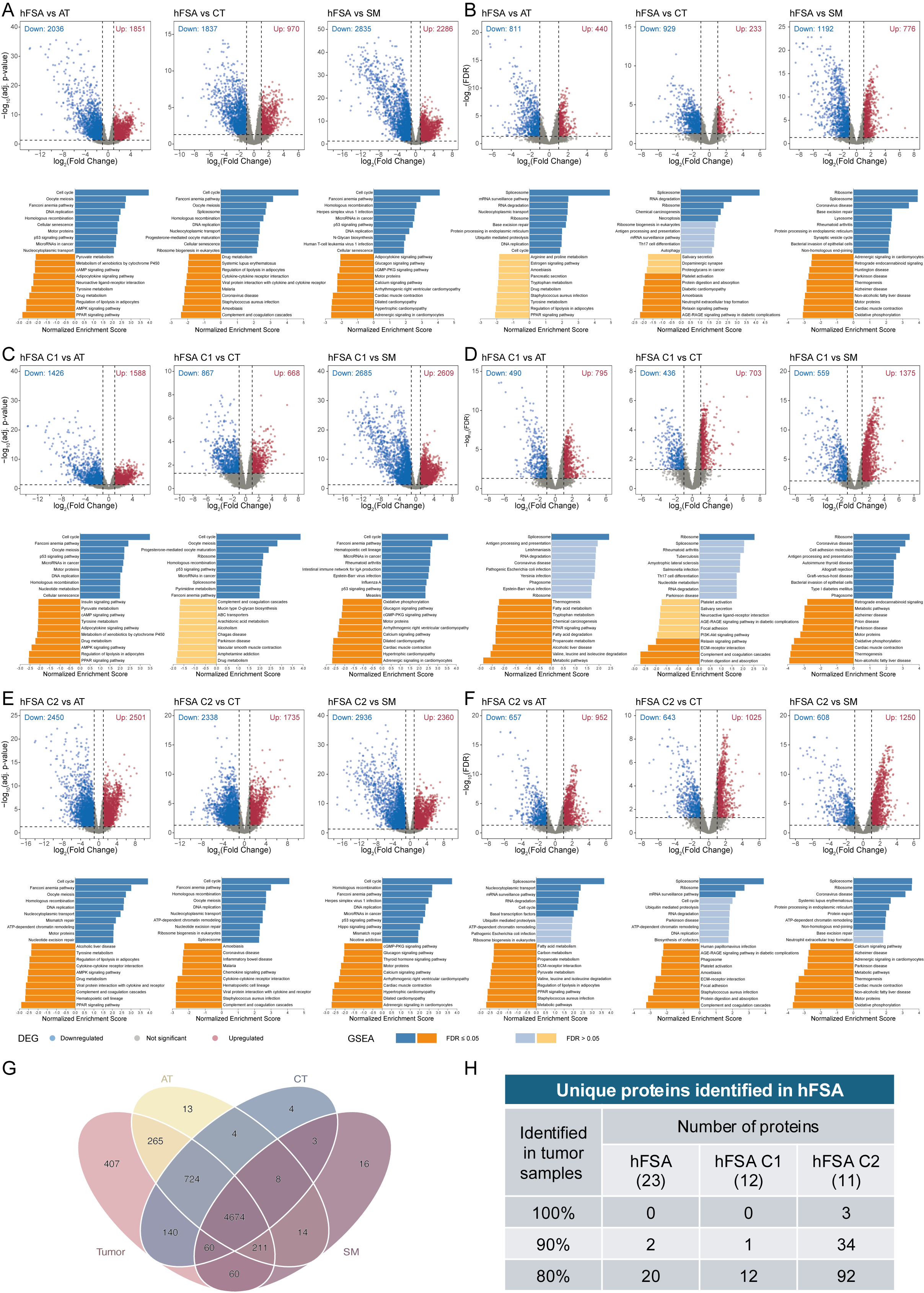

Supplementary figure 16. Characterization of hFSA compared to NT.

A. Differentially expressed genes (|log_2_(FC)| ≥1 & adj. p-value <0.05) in hFSA compared to AT, CT and SM with subsequent GSEA using Gene Ontology terms of Biological Processes.
B. Differentially expressed proteins (|log_2_(FC)| ≥1 & FDR <0.05) in hFSA compared to AT, CT and SM with subsequent GSEA using KEGG pathways.

C+D.	Differentially expressed C) genes and D) proteins (|log_2_(FC)| ≥1 & adj. p-value/FDR <0.05) in hFSA C1 compared to AT, CT and SM with subsequent GSEA using KEGG pathways.

E+F.	Differentially expressed C) genes and D) proteins (|log_2_(FC)| ≥1 & adj. p-value/FDR <0.05) in hFSA C1 compared to AT, CT and SM with subsequent GSEA using KEGG pathways.

G.	Venn diagram showing proteins identified per tissue, with 407 proteins uniquely detected in hFSA tumour tissue but not in the normal tissue.

H. Table highlighting the number of tumour unique proteins identified in G which are detected in more than 80, 90 and 100% of tumour samples. 20 proteins were exclusively detected in hFSA tumour tissue in ≥80%, and 2 proteins in even ≥90% of all samples. Separating hFSA into C1 and C2, revealed even 92 proteins in ≥80%, and 34 proteins in ≥90% uniquely expressed in tumours belonging to hFSA C2.

Supplementary Figure 17.
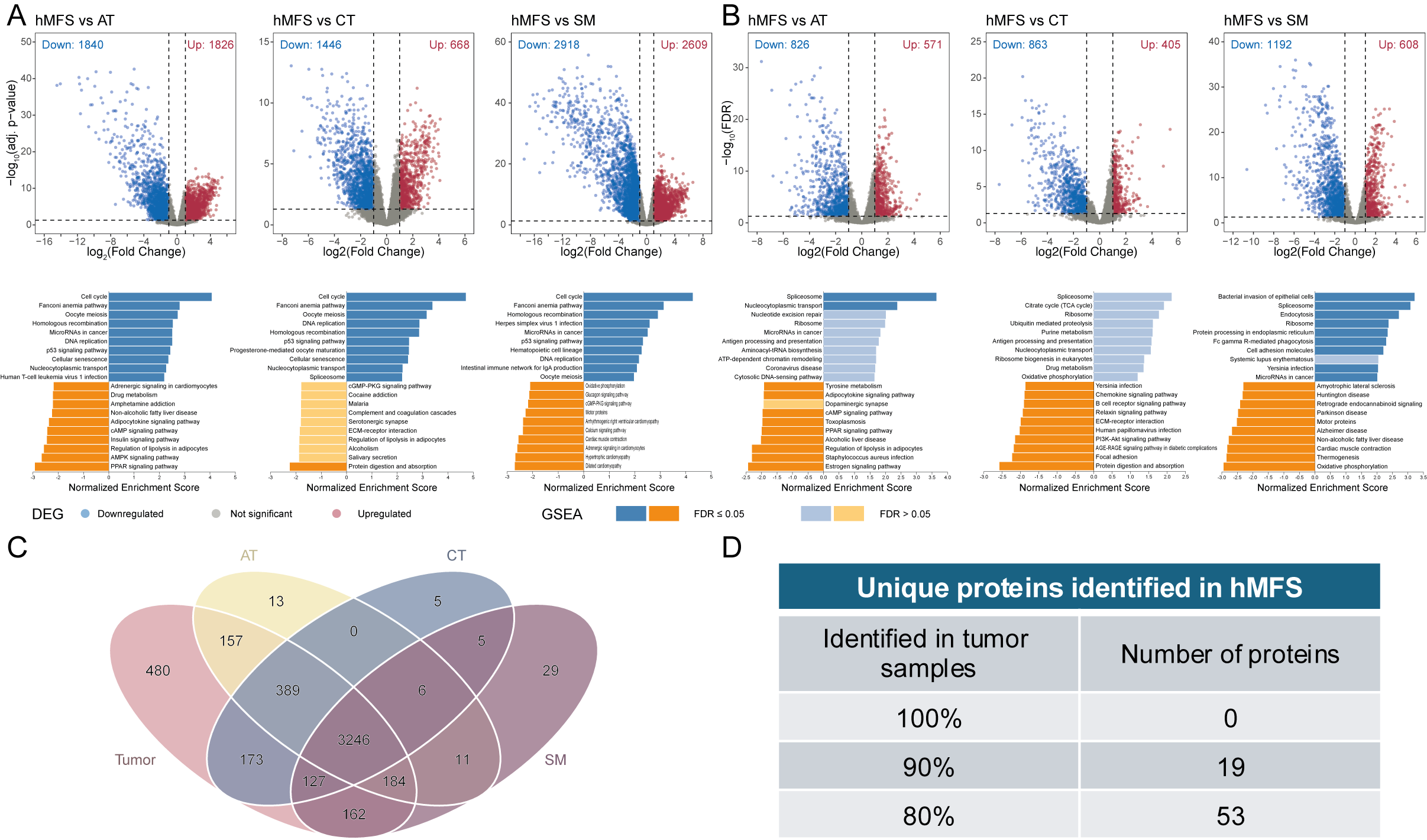

Supplementary figure 17. Characterization of hMFS compared to NT.

A. Differentially expressed genes (|log_2_(FC)| ≥1 & adj. p-value <0.05) in hMFS compared to AT, CT and SM with subsequent GSEA using KEGG pathways.
B. Differentially expressed proteins (|log_2_(FC)| ≥1 & FDR <0.05) in hMFS compared to AT, CT and SM with subsequent GSEA using KEGG pathways.
C. Venn diagram showing proteins identified per tissue, with 480 proteins uniquely detected in hMFS tumour tissue.
D. Table highlighting 53 proteins exclusively detected in hMFS tumour tissue in ≥80%, and 19 proteins in even ≥90% of all samples.

Supplementary Figure 18.

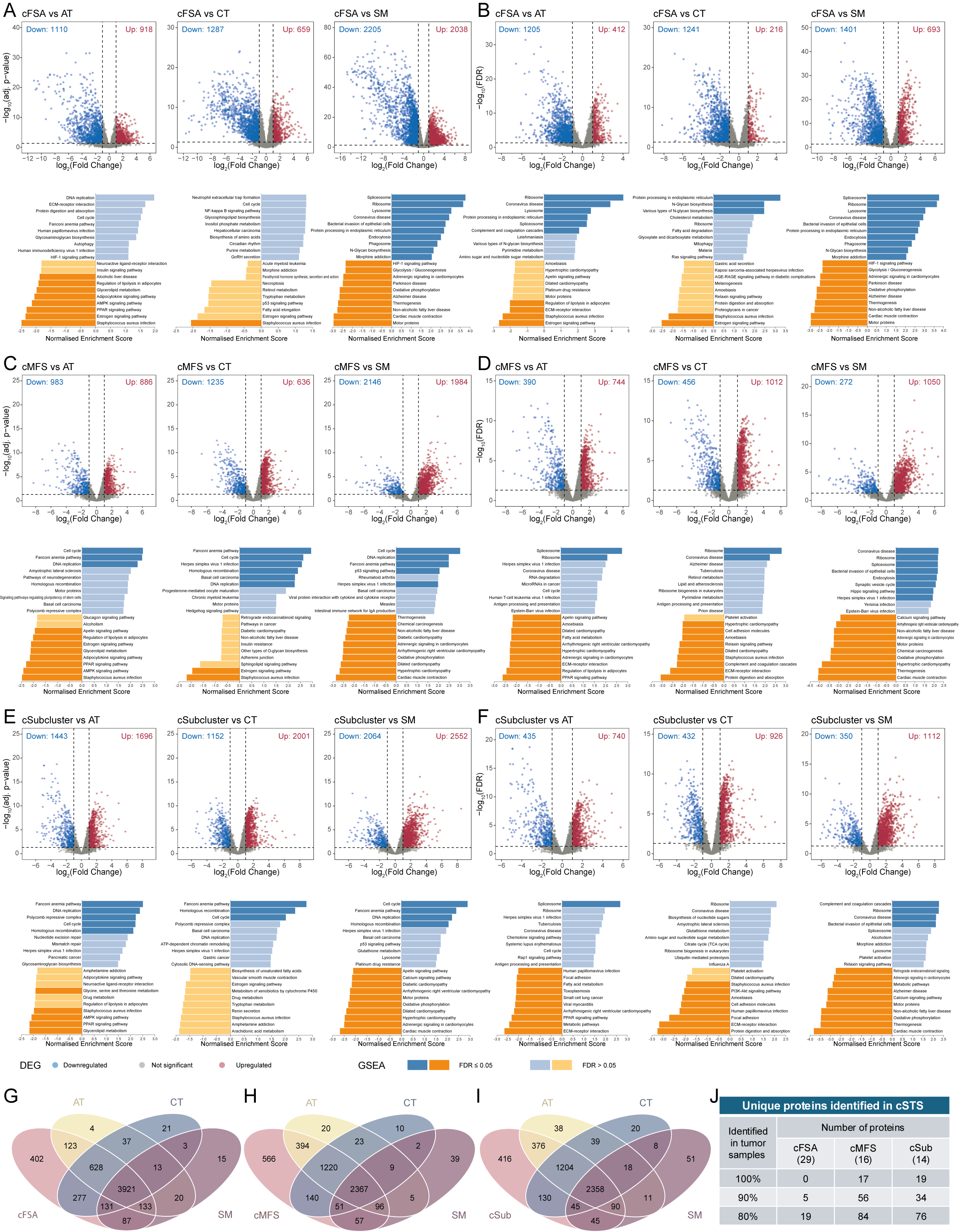

Supplementary figure 18. Characterization of canine STS entities compared to NT.

A+B.	Differentially expressed A) genes and B) proteins (|log_2_(FC)| ≥1 & adj. p-value/FDR <0.05) in cFSA compared to AT, CT and SM with subsequent GSEA using Gene Ontology terms of Biological Processes.

C+D.	Differentially expressed C) genes and D) proteins (|log_2_(FC)| ≥1 & adj. p-value/FDR <0.05) in cMFS compared to AT, CT and SM with subsequent GSEA using KEGG pathways.

E+F.	Differentially expressed C) genes and D) proteins (|log_2_(FC)| ≥1 & adj. p-value/FDR <0.05) in cSubcluster compared to AT, CT and SM with subsequent GSEA using KEGG pathways.

G+H+I.)	Venn diagram showing proteins identified per tissue detected in tumour tissue of G) cFSA, H) cMFS and I) cSubcluster.

J. Table highlighting proteins exclusively detected in tumour tissue with more than ≥80, ≥90% or all samples of cFSA, cMFS or cSubcluster.

Supplementary Figure 19

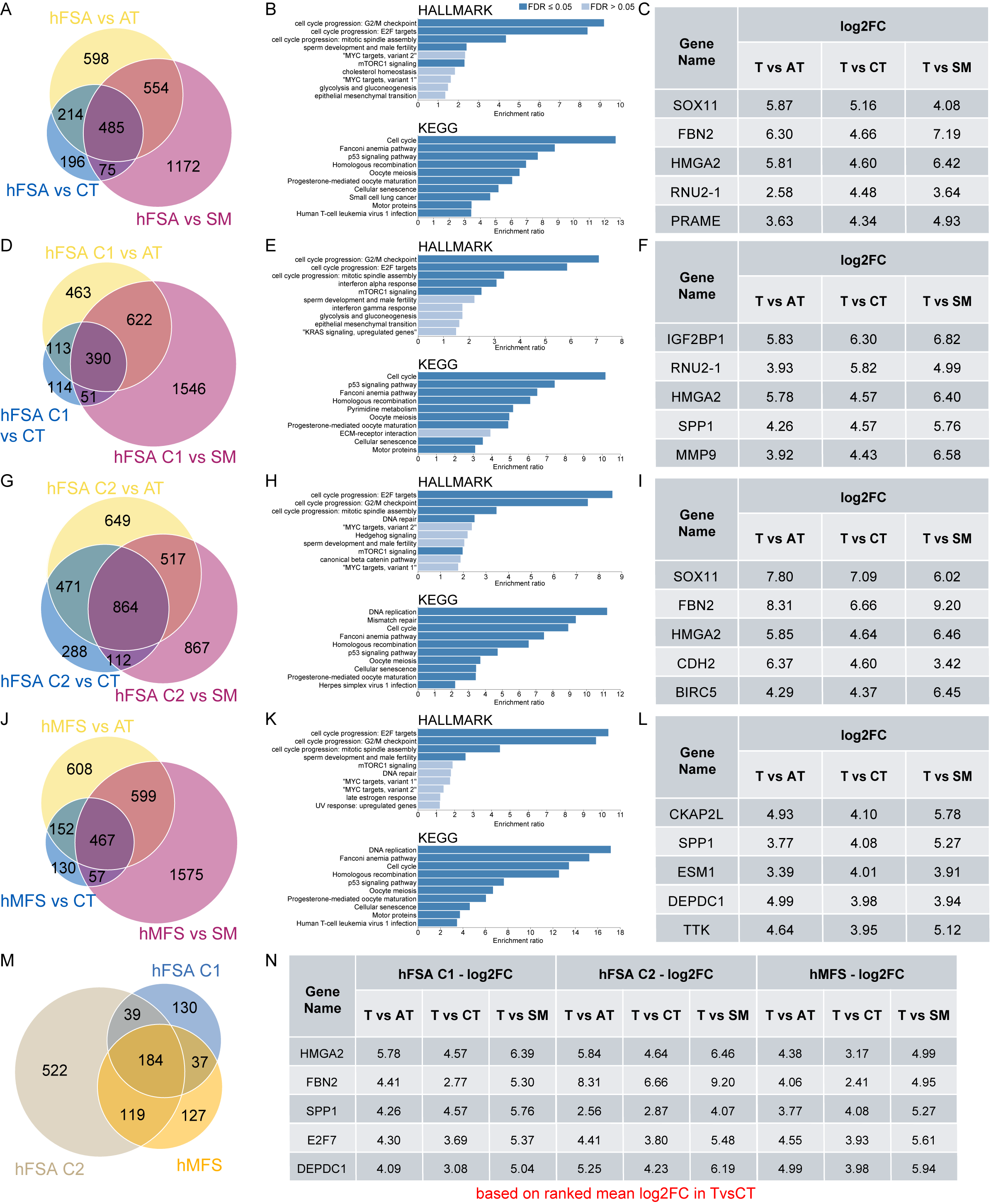

Supplementary Figure 19. Differential gene expression of human tumour clusters.

A – C)	Differential gene expression of hFSA (whole cohort) showing A) Venn diagram with overlapping number of identified targets (log_2_(FC) > 1 & adj. p-value < 0.05) compared to NT, B) GSEA analysis of 485 targets commonly upregulated in tumour versus NT, and C) top5 targets ranked by log_2_(FC) in hFSA vs CT.

D – F)	Differential gene expression of hFSA C1 showing D) Venn diagram with overlapping number of identified targets (log_2_(FC) > 1 & adj. p-value < 0.05) compared to NT, E) GSEA analysis of 390 targets commonly upregulated in tumour versus NT, and F) top5 targets ranked by log_2_(FC) in hFSA C1 vs CT.

G – H)	Differential gene expression of hFSA C2 showing D) Venn diagram with overlapping number of identified targets (log_2_(FC) > 1 & adj. p-value < 0.05) compared to NT, E) GSEA analysis of 864 targets commonly upregulated in tumour versus NT, and F) top5 targets ranked by log_2_(FC) in hFSA C2 vs CT.

J – L)	Differential gene expression of hMFS showing D) Venn diagram with overlapping number of identified targets (log_2_(FC) > 1 & adj. p-value < 0.05) compared to NT, E) GSEA analysis of 467 targets commonly upregulated in tumour versus NT, and F) top5 targets ranked by log_2_(FC) in hMFS vs CT.

M)	Venn diagram showing overlap of interesting targets between hFSA C1, hFSA C2 and hMFS.

N)	Top5 common targets ranked by mean log_2_(FC) in tumour vs CT.

Supplementary Figure 20.

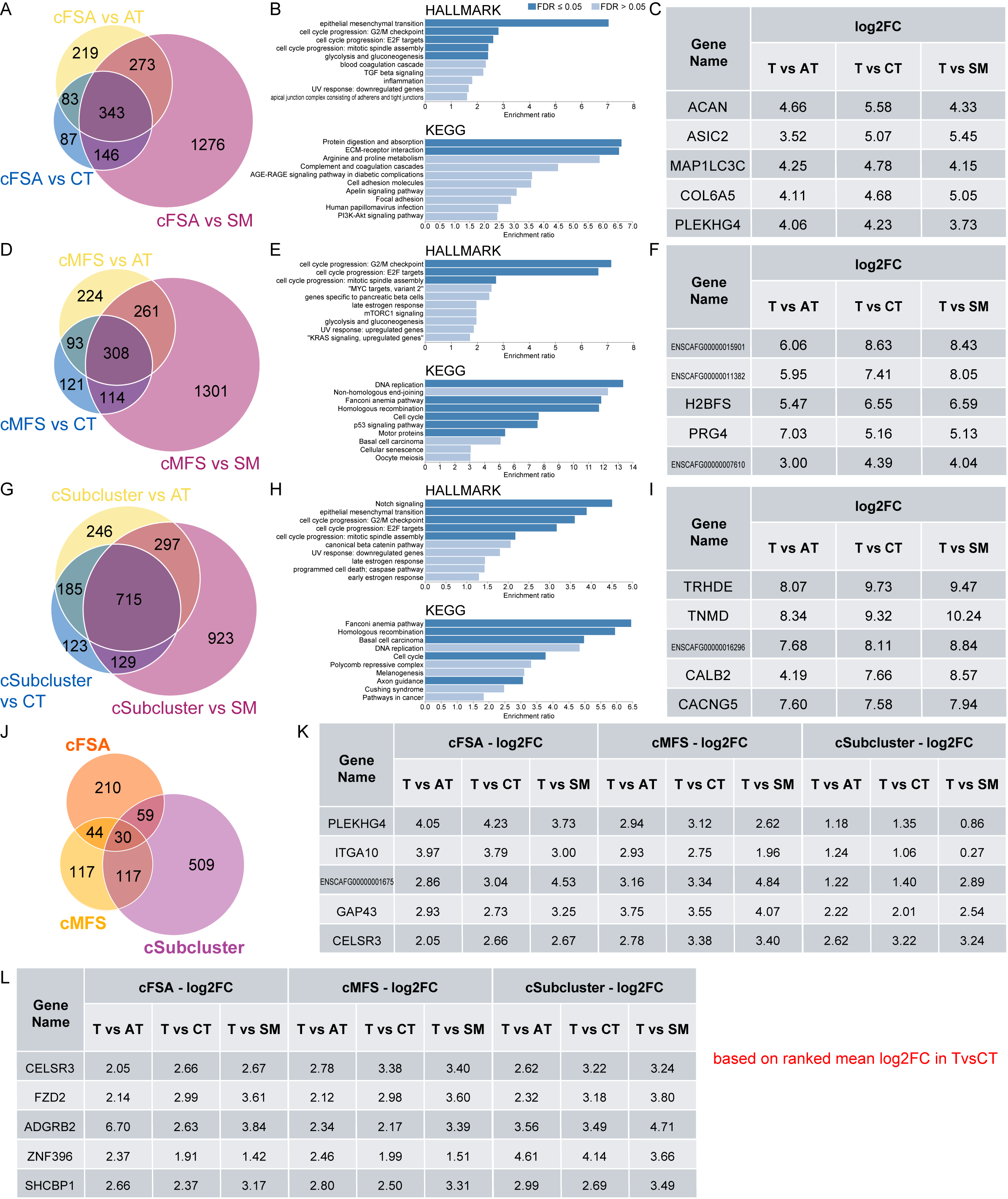

Supplementary Figure 20. Differential gene expression of canine tumour clusters.

A – C)	Differential gene expression of cFSA showing A) Venn diagram with overlapping number of identified targets (log_2_(FC) > 1 & adj. p-value < 0.05) compared to NT, B) GSEA analysis of 343 targets commonly upregulated in tumour versus NT, and C) top5 targets ranked by log_2_(FC) in cFSA vs CT.

D – F)	Differential gene expression of cMFS showing D) Venn diagram with overlapping number of identified targets (log_2_(FC) > 1 & adj. p-value < 0.05) compared to NT, E) GSEA analysis of 308 targets commonly upregulated in tumour versus NT, and F) top5 targets ranked by log_2_(FC) in cMFS vs CT.

G – H)	Differential gene expression of cSubcluster showing D) Venn diagram with overlapping number of identified targets (log_2_(FC) > 1 & adj. p-value < 0.05) compared to NT, E) GSEA analysis of 715 targets commonly upregulated in tumour versus NT, and F) top5 targets ranked by log_2_(FC) in cSubcluster vs CT.

J)	Venn diagram showing overlap of interesting targets between cFSA, cMFS and cSubcluster.

K)	Top5 common targets ranked by mean log_2_(FC) in tumour vs CT between cFSA and cMFS.

Supplementary Figure 21

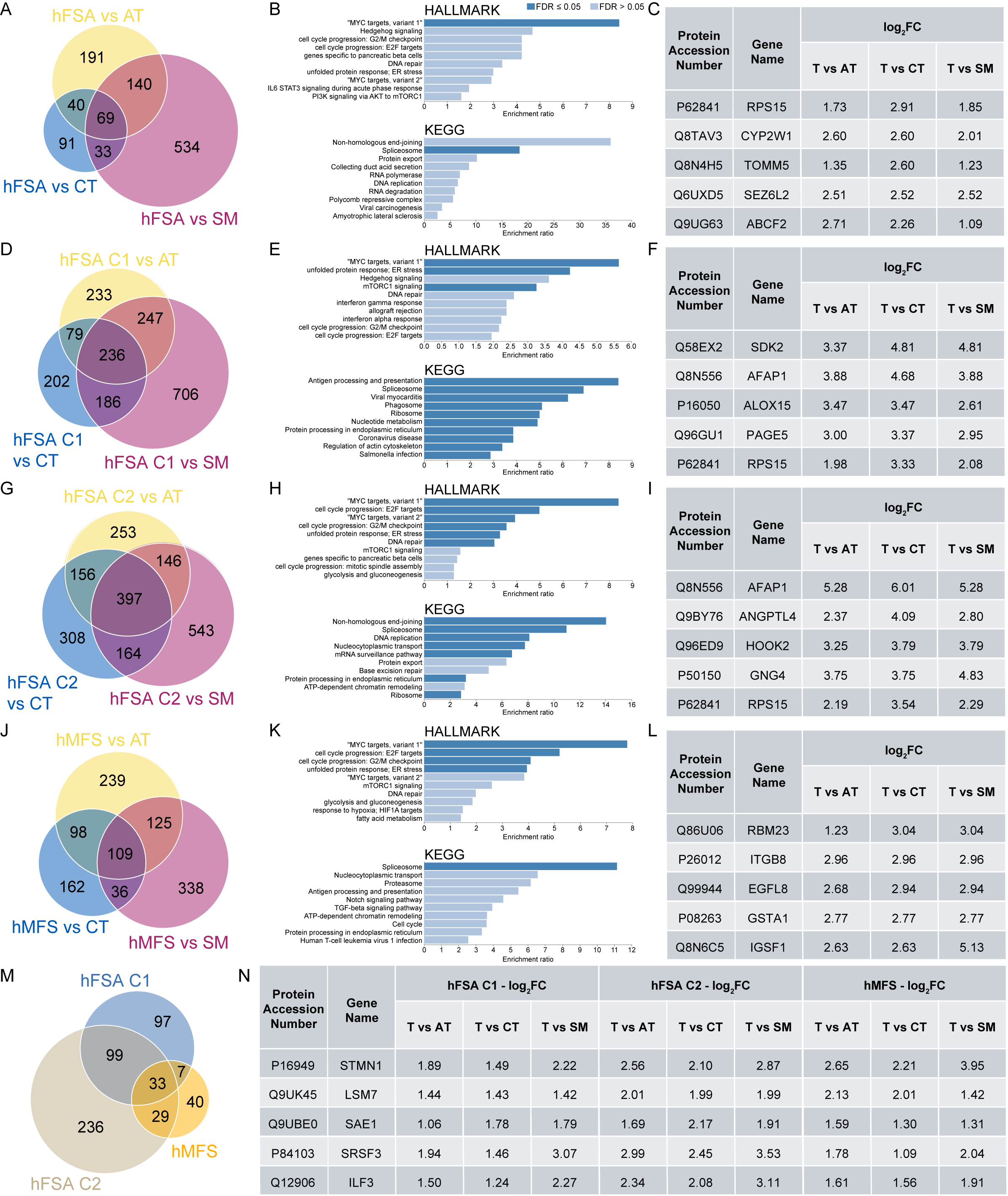

Supplementary Figure 21. Differential protein expression of human tumour clusters.

A – C)	Differential protein expression of hFSA (whole cohort) showing A) Venn diagram with overlapping number of identified targets (log_2_(FC) > 1 & adj. FDR < 0.05) compared to NT, B) GSEA analysis of 69 targets commonly upregulated in tumour versus NT, and C) top5 targets ranked by log_2_(FC) in hFSA vs CT.

D – F)	Differential gene expression of hFSA C1 showing D) Venn diagram with overlapping number of identified targets (log_2_(FC) > 1 & adj. FDR < 0.05) compared to NT, E) GSEA analysis of 236 targets commonly upregulated in tumour versus NT, and F) top5 targets ranked by log_2_(FC) in hFSA C1 vs CT.

G – H)	Differential protein expression of hFSA C2 showing D) Venn diagram with overlapping number of identified targets (log_2_(FC) > 1 & adj. FDR < 0.05) compared to NT, E) GSEA analysis of 397 targets commonly upregulated in tumour versus NT, and F) top5 targets ranked by log_2_(FC) in hFSA C2 vs CT.

J – L)	Differential protein expression of hMFS showing D) Venn diagram with overlapping number of identified targets (log_2_(FC) > 1 & adj. FDR < 0.05) compared to NT, E) GSEA analysis of 109 targets commonly upregulated in tumour versus NT, and F) top5 targets ranked by log_2_(FC) in hMFS vs CT.

M)	Venn diagram showing overlap of interesting targets between hFSA C1, hFSA C2 and hMFS.

N)	Top5 common targets ranked by mean log_2_(FC) in tumour vs CT.

Supplementary Figure 22.

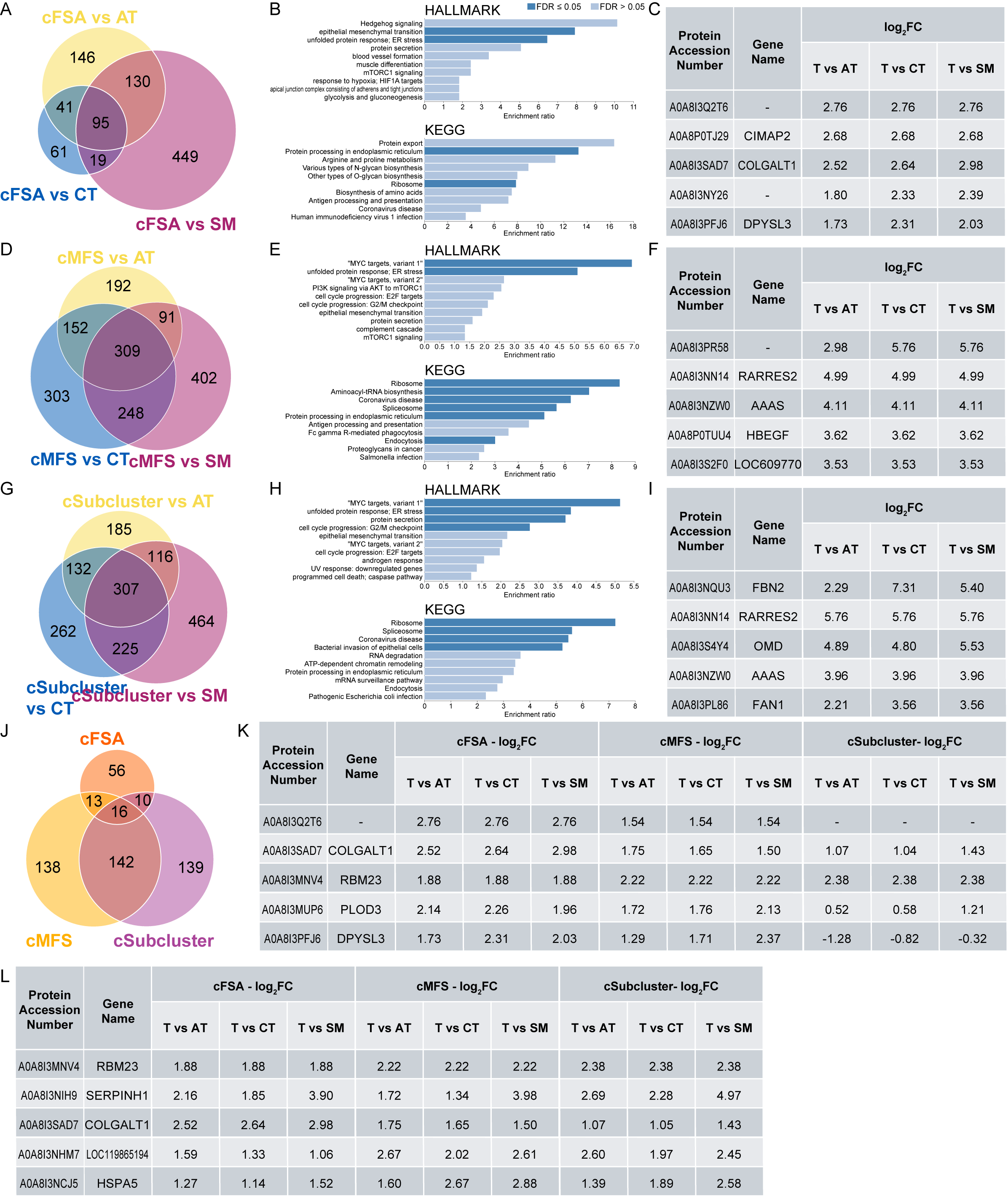

Supplementary Figure 22. Differential protein expression of canine tumour clusters.

A – C) Differential protein expression of cFSA showing A) Venn diagram with overlapping number of identified targets (log_2_(FC) > 1 & adj. p-value < 0.05) compared to NT, B) GSEA analysis of 95 targets commonly upregulated in tumour versus NT, and C) top5 targets ranked by log_2_(FC) in cFSA vs CT.

D – F) Differential protein expression of cMFS showing D) Venn diagram with overlapping number of identified targets (log_2_(FC) > 1 & adj. p-value < 0.05) compared to NT, E) GSEA analysis of 309 targets commonly upregulated in tumour versus NT, and F) top5 targets ranked by log_2_(FC) in cMFS vs CT.

G – H) Differential protein expression of cSubcluster showing D) Venn diagram with overlapping number of identified targets (log_2_(FC) > 1 & adj. p-value < 0.05) compared to NT, E) GSEA analysis of 307 targets commonly upregulated in tumour versus NT, and F) top5 targets ranked by log_2_(FC) in cSubcluster vs CT.

J)	Venn diagram showing overlap of interesting targets between cFSA, cMFS and cSubcluster.

K)	Top5 common targets ranked by mean log_2_(FC) in tumour vs CT between cFSA and cMFS.
